# Purified fibers in chemically defined synthetic diets destabilize the gut microbiome of an omnivorous insect model

**DOI:** 10.1101/2024.05.15.594388

**Authors:** Rachel L. Dockman, Elizabeth A. Ottesen

## Abstract

The macronutrient composition of a host’s diet shapes its gut microbial community, with dietary fiber in particular escaping host digestion to serve as a potent carbon source for gut microbiota. Despite widespread recognition of fiber’s importance to microbiome health, nutritional research often fails to differentiate hyper-processed fibers from cell-matrix derived intrinsic fibers, limiting our understanding of how individual polysaccharides influence the gut community. We use the American cockroach (*Periplaneta americana*) as a model system to dissect the response of complex gut microbial communities to diet modifications that are impossible to test in traditional host models. Here, we designed synthetic diets from lab-grade, purified ingredients to identify how the cockroach microbiome responds to six different carbohydrates (chitin, methylcellulose, microcrystalline cellulose, pectin, starch, xylan) in otherwise balanced diets. We show via 16S rRNA gene profiling that these synthetic diets reduce bacterial diversity and alter the phylogenetic composition of cockroach gut microbiota in a fiber-dependent manner, regardless of the vitamin and protein content of the diet. Comparisons with cockroaches fed whole-food diets reveal that synthetic diets induce blooms in common cockroach-associated taxa and subsequently fragment previously stable microbial correlation networks. Our research leverages an unconventional microbiome model system and customizable lab-grade artificial diets to shed light on how purified polysaccharides, as opposed to nutritionally complex intrinsic fibers, exert substantial influence over a normally stable gut community.

## 2 Introduction

The gut microbiome is a key player in host metabolism and homeostasis; it extracts energy from recalcitrant dietary components, provisions essential nutrients, and stimulates the host’s immune system to protect against pathogens and toxins (Bäckhed et al., 2005; Huttenhower et al., 2012; Thaiss et al., 2016; Belkaid and Harrison, 2017). These benefits to the host are contingent upon the microbiota present, which themselves are selected through external pressure such as host genetics environment, and diet (Goodrich et al., 2014; Sonnenburg et al., 2016; Kurilshikov et al., 2017; Greene et al., 2020; Kurilshikov et al., 2021). Diet has gained particular attention as the most easily manipulated of these factors, and a clear relationship exists between microbially derived metabolic products from the gut microbiome and overall host health (Tanes et al., 2021; Wolter et al., 2021).

Shifts in the ratios and sources of metabolizable macronutrients (fats, carbohydrates, protein) are frequently identified as drivers of diet-associated microbiota alterations, but the most important component to resident gut bacteria is what bypasses host digestion relatively untouched: fiber (Walker et al., 2011; Rastall et al., 2022). Dietary fiber consists of plant-derived structural carbohydrates that most animals are unable to process and are thus key to maintaining a diverse, beneficial gut microbial community. However, performing research relating dietary fiber consumption to gut microbiota within a host organism presents several challenges. Whole foods contain “intrinsic fibers”, an assortment of carbohydrates characterized by source-specific molecular structures that form close associations with plant proteins and cell matrix components (Tuncil et al., 2020; Puhlmann and de Vos, 2022). These more labile components can obscure the influence of individual structural polysaccharides on the gut community, especially considering the high diversity of carbohydrate degrading machinery found across individual lineages of gut microbiota (Kaur et al., 2019; Liu et al., 2020; Villa et al., 2020). Purified fibers present an alternative that control for these variable compounds, but mammalian models have complex nutritional needs that limit the extent of dietary manipulation possible before introducing host stresses. As a result, *in vivo* dietary research frequently requires that pure fibers be used as dietary supplements rather than primary carbohydrate source, obscuring microbial responses to the fiber itself without preventing alternative energy sources from being prioritized by gut bacteria. Invertebrate models offer more flexibility, but well-known insect models such as *Drosophila* have limited dietary range that poorly reflect the community dynamics found in mammalian host species (Lee and Brey, 2013; Lesperance and Broderick, 2020). To address this challenge, we are developing the omnivorous American cockroach (*Periplaneta americana*) as a model of microbiome dynamics that extends our understanding of human-relevant bacteria while leveraging the benefits of invertebrate research.

Omnivorous cockroaches such as *P. americana* are colonized by a complex hindgut microbiome that is taxonomically similar to the human colonic flora, consisting of many shared microbial lineages such as Bacteroidota, Firmicutes (now Bacillota), and Proteobacteria (now Pseudomonadota) (Cruden and Markovetz, 1987; Schauer et al., 2012; Tinker and Ottesen, 2016). Additionally, cockroach gut microbiota play functionally analogous roles in host nutrition, with hindgut bacteria fermenting otherwise indigestible dietary components into volatile fatty acids that can be utilized by the host as an energy supplement or influence host development (Zurek and Keddie, 1996; Ayayee et al., 2018; Jahnes and Sabree, 2020; Vera-Ponce de León et al., 2021). Further, the extreme dietary flexibility conferred by *Blattabacterium* endosymbionts and the unique ability for cockroaches to store nitrogen as uric acid for future protein synthesis facilitates extensive dietary manipulation (Cochran et al., 1979; Sabree et al., 2009; Ayayee et al., 2016). Studies of cockroach gut microbiome responses to diet have generated contrasting responses, with multiple large-scale studies finding that the gut microbiome is highly stable following dietary shifts (Schauer et al., 2014; Tinker and Ottesen, 2016; Lampert et al., 2019), while others have demonstrated shifts in response to select diets (Bertino-Grimaldi et al., 2013; Pérez-Cobas et al., 2015; Zhu et al., 2022). Currently, there is no consensus on why these studies produced differing results and comparison is difficult due to inconsistent use of synthetic, natural, or whole food diets across studies. Whole food or natural diets may obscure bacterial responses to nutritional contents given their complexity, while synthetic diets are more amenable to precise dietary changes, thus controlling for more variables (Okarter and Liu, 2010; Koropatkin et al., 2012; Adesogan et al., 2014; Tuncil et al., 2017). Artificial diets have been successfully developed and used as replacement for natural diets in insects with far more specialized dietary needs than cockroaches, implying that cockroaches are an ideal candidate for use with lab-synthesized diets (Piper et al., 2014; Talyuli et al., 2015; Gonzales et al., 2018; Majumder et al., 2020).

To facilitate precise manipulation of dietary composition in cockroaches, we have developed a series of synthetic cockroach diets based on the work of early entomologists (Noland et al., 1949; Haydak, 1953; Hamilton and Schal, 1988). These artificial diets serve as a nutritionally complete base to isolate the influence of specific dietary components on the *P. americana* hindgut microbiome, a community known to be resistant to dietary manipulation when fed macronutrient-biased whole food diets (Tinker and Ottesen, 2016). Using these synthetic diets as a base, we tested a spectrum of polysaccharides as the primary carbon and energy source to identify if the hindgut microbiome responds to specific fibers without obfuscation by intrinsic fiber components. We found that these diets resulted in much stronger impacts on gut microbiome composition than highly divergent whole food diets, with long-chain polysaccharide source exerting the largest effect even when diets are modified in their protein and micronutrient composition. Our work will facilitate future studies of gut microbiome responses to fine-scale dietary composition in the cockroach and shed light on how hyper-processed synthetic diets, which superficially appear to be nutritionally complete, destabilize a complex gut microbiome.

## 3 Materials and Methods

### 3.1 Insects and Experimental Conditions

Our *Periplaneta americana* colony has been maintained in captivity at the University of Georgia for over a decade. Mixed age and sex stock insects are maintained at room temperature in glass aquarium tanks with wood chip bedding and cardboard tubes for shelter in a 12:12 light:dark cycle. Water via cellulose sponge fit to a Tupperware reservoir and dog chow (Purina ONE chicken & rice formula, approximately 26% protein, 16% fat, and 3% fiber) are provided to stock colonies *ad libitum*.

### 3.2 Synthetic Diets

The synthetic diets created for dietary testing were designed to provide balanced nutrition while remaining malleable to component manipulation. Diets contained Vanderzant vitamin mix (MP Biomedicals, Irvine, CA, USA), Wesson salt mix (MP Biomedicals), peptone (Amresco, VWR International, Radnor, PA, USA), casein (Sigma-Aldrich, St. Louis, MO, USA), and cholesterol (VWR); amounts are listed in **Table 1**. The dry ingredients were suspended in sufficient volumes of diH_2_O to create a batter or dough, formed into pellets, then dehydrated at 65°C until they were sufficiently dry to maintain shape. Food pellets were stored at −20°C until use.

**Table 1.** Synthetic diet compositions. CHO: carbohydrate; MeC: methylcellulose; MCC: microcrystalline cellulose; P-: protein deficient; V-: vitamin deficient; *: canned tuna was dried prior to weighing.

| Diet Type | CHO % | Casein % | Peptone % | Mineral Mix % | Vitamin Mix % | Cholesterol % | Diet/CHO |
| --- | --- | --- | --- | --- | --- | --- | --- |
| Standard: Polysaccharide | 70.5 | 17 | 8 | 3 | 0.5 | 1 | Chitin, MeC, MCC, Pectin, Starch, Xylan |
| Standard: Simple Sugar | 70.5 | 17 | 8 | 3 | 0.5 | 1 | Cellobiose, Glucose, Xylose |
| Protein Deficient | 95.5 | 0 | 0 | 3 | 0.5 | 1 | MCC P-, Xylan P- |
| Vitamin Deficient | 72 | 17 | 8 | 3 | 0 | 0 | MCC V-, Xylan V- |
| Tuna-Amended | 70.5 | 25% tuna |  | 3 | 0.5 | 1 | MCC, Xylan |

In most experiments, the only component changed was the carbohydrate source. Polysaccharides used include microcrystalline cellulose (MCC; Sigma-Aldrich), methylcellulose (Sigma-Aldrich), xylan from corn core (TCI Chemicals, Portland, OR, USA), pectin from apple (Sigma-Aldrich), starch (Sigma-Aldrich), and chitin (Alfa Aesar, Ward Hill, MA, USA). For simple sugar diet variations, cellobiose (Sigma-Aldrich), glucose (Sigma-Aldrich), and xylose (Acros Organics, VWR International) were used as the carbohydrate component.

### 3.3 Experimental Design

Experimental conditions were prepared as described in (Tinker and Ottesen, 2016). Briefly, mixed-sex healthy adult insects were transferred from the stock colony to plastic tanks containing pebbles and bleached polyvinyl chloride tubes for footing and shelter, respectively. Food and water were provided *ad libitum* in rigid plastic or glass dishes following two days of food restriction and habituation. Dietary treatment lasted two weeks, during which debris, oothecae, and lethargic insects were removed daily.

Upon completion of dietary treatments, each insect was isolated in a sterile culture plate and placed on ice until torpid. Sternites were removed with sterile forceps to expose the intact gut and fat body tissue was cleaned away. The cleaned gut was frozen on a sterile aluminum dish on dry ice and divided into foregut, midgut, and hindgut sections for collection in 500-800µL phosphate-buffered saline (1X PBS). Gut contents and tissue-attached bacteria were disrupted with a sterile pestle, and the samples stored at –20°C until DNA extraction. For this study, only the hindgut community was analyzed due to higher microbial density and activity than in other gut regions.

### 3.4 DNA Extraction

DNA was extracted from 200µL aliquots of individual samples using the EZNA Bacterial DNA Kit (Omega Biotek, Norcross, GA, USA) with some modifications. Sample aliquots were centrifuged at 5000g for 10min, with the resulting pellet resuspended in 100µL TE buffer plus 10µL lysozyme (50mg/mL) and incubated for 30 min at 37°C. Following incubation, samples were vortexed with glass beads (25mg, Omega Biotek) for 5 min at 3000rpm, then incubated at 55°C for one hour with 100µL TL buffer, 20µL proteinase K and continuous 600rpm shaking. The kit protocol was followed for additional incubations with BL buffer and DNA isolation using the provided column. DNA was eluted into 50µL of provided Elution Buffer and quantified using either a Nanodrop Lite spectrophotometer (Thermo Scientific) or the Take3 plate for BioTek plate readers (Agilent).

### 3.5 16S rRNA Gene Library Preparation and Sequencing

The V4 region of the 16S rRNA gene was amplified via 2-step polymerase chain reaction (PCR) from individual hindgut lumen samples as previously described in (Tinker and Ottesen, 2016; 2020; 2021). Both PCR reactions used 0.02U/L Q5 Hot Start high-fidelity DNA polymerase (New England BioLabs, Ipswich, MA, USA) with 200M dNTPs and 0.5M forward and reverse primers in 1M Q5 reaction buffer. The first 10µL reaction containing 3ng DNA and primers targeting the V4 region (**515F**: GTGCCAGCMGCCGCGGTAA; **806R**: GGACTACHVGGGTWTCTAAT) was performed under the following conditions: activation at 98°C for 30s; 15 cycles of 98°C for 10s, 52°C for 30s, and 72°C for 30s; final extension at 72°C for 2 min. Immediately following amplification, 9µL of the first reaction was added to 21µL of Q5 reaction mix containing barcoded primers with adaptor sequences for Illumina sequencing (Caporaso et al., 2011). Cycling was performed as follows: activation at 98°C for 30s; 4 cycles of 98°C for 10s, 52°C for 10s, and 72°C for 30s; 6 cycles of 98°C for 10s and 72°C for 1 min; final extension at 72°C for 2 min.

After product size verification via gel electrophoresis, samples were cleaned as instructed in Omega Biotek’s Cycle Pure kit, quantified, and pooled for equimolar representation of each sample. Prepared libraries were sent to the Georgia Genomics and Bioinformatics Core at the University of Georgia for 250 base pair paired-end Illumina MiSeq sequencing.

### 3.6 Amplicon Sequence Variant Generation

Each dataset collected in this study was processed separately in R (version 4.2.1) by sequencing run using R package DADA2 (version 1.24.0), with the cumulative Amplicon Sequence Variants (ASVs) generated input as a priors table for each successive run (Callahan et al., 2016; RStudioTeam, 2020). To allow for comparison with this dataset, raw data from previous research in the Ottesen lab were reprocessed to generate ASVs following the same procedures as in this current study (Tinker and Ottesen, 2016). All sequence tables produced by these datasets were combined by ASV sequence prior to taxonomy assignment to ensure continuity in naming. Taxonomy was assigned using DADA2 and the ARB Silva v138 classifier to the species level, uniquely numbered, and filtered to remove sequences matching eukaryotic (chloroplast, mitochondria) or endosymbiotic *Blattabacterium* DNA (Quast et al., 2013; Callahan et al., 2016).

### 3.7 Community Analysis

Alpha and beta diversity analyses were performed via the R package vegan (version 2.6-4) (Jari Oksanen and Peter Solymos, 2019). Samples were rarefied prior to diversity analysis to 7924 reads for comparisons between synthetic and/or whole food diets, 9685 reads for analysis of repeat xylan and MCC diet experiments, and 12274 reads for follow-up experiments exploring nutrient deficiencies and simple sugar carbohydrates. Alpha diversity was measured via Shannon index, the count of ASVs observed in rarefied samples, and Pielou’s evenness (calculated as Shannon/log(Observed)). Weighted and unweighted Bray-Curtis dissimilarities were calculated, assessed for dispersion, and plotted using the vegan functions vegdist, betadisper, and metaMDS. Statistics for alpha diversity indices were calculated with the Wilcoxon rank sum test (pairwise comparisons) and Kruskal-Wallis test (multi-group comparisons). The significance of community composition differences observed in beta diversity measures was assessed using PERMANOVA (adonis2 in vegan package). Beta dispersion was further examined through the Tukey’s HSD test for pairwise comparisons and ANOVA for multi-group comparisons.

Differential abundance analysis was conducted using DESeq2 (version 1.36) (Love et al., 2014). For identification of ‘diet-characteristic taxa”, raw count data for the synthetic diet set (n=66) was filtered to exclude ASVs present in less than 5 samples and run through the ‘DESeq’ command with parameters ‘fitType = “local”’ and ‘design = ∼ Diet’. Pairwise result tables were obtained for all diet comparisons and filtered for significant data, defined as having an adjusted p-value smaller than 0.05 and a baseMean larger than 10. ASVs significantly upregulated for one diet vs the other five diets were identified as diet-characteristic (n=76) and used to generate the heatmap in **Supplement 3**. For comparison of “synthetic vs whole”, raw count data for both diet sets (n=125) were combined and filtered to exclude ASVs that appeared in fewer than 5 samples. DESeq was run with parameters ‘fitType = “local”’ and ‘design = ∼ Diet_Type’ to identify differentially abundant ASVs between the diet types. The resulting baseMean and log2 fold change were used to generate the MA plots in **Figure 4**.

UpSetR (version 1.4) was employed to visualize intersecting sets of taxa, providing insights into the distribution of taxonomic features across samples (Conway et al., 2017). For UpSet analysis, samples were rarefied to 7924 reads then collapsed together to obtain total counts per diet. Both a presence/absence table and a proportion table were generated from these data, with the presence/absence table used for UpSet graph generation. Relative abundance of each set was calculated using the proportion table with ASVs collapsed per set and visualized as pie charts within the UpSet graph.

Co-correlation analysis was conducted to evaluate the impact of synthetic diets on microbial interaction networks using the SparCC procedure (Friedman and Alm, 2012). Networks were constructed separately for the synthetic diet group and the whole food diet group using sequence count tables that were filtered to only include ASVs with at least 5 representatives present in 25% of samples (synthetic: 17 samples; whole food: 15 samples), preventing spurious correlations from rare taxa. SparCC was implemented in R with standard parameters, and the resultant networks were characterized and analyzed with the igraph R package (version 1.5.1) (Csardi and Nepusz, 2006; Csárdi et al., 2024). Networks were pruned to contain only edges with a correlation absolute value of at least 0.4 and exported into Cytoscape for visualization using the edge-weighted spring embedded layout method (Shannon et al., 2003).

## 4 Results

### 4.1 Impacts of synthetic diets on gut microbiome diversity and community composition

We formulated a series of synthetic diets composed of a fixed base of 25% protein amended with dietary salts, vitamins, and cholesterol while differing only in complex carbohydrate type. Initial experiments utilized five alternative polysaccharide sources: chitin, methylcellulose, microcrystalline cellulose (MCC), pectin, or xylan. Following initial analysis of these results, we tested an additional starch-based diet. The prepared diets were readily consumed by the cockroaches in all cases.

To evaluate the impact of these diets on the gut community, each diet was fed to adult cockroaches for a period of 14 consecutive days, after which the insects were sacrificed, and their hindgut dissected out for 16S rRNA gene library sequencing. Following library preparation and sequencing, we used DADA2 to obtain 2,321,848 quality-controlled, assembled sequences assigned to 3308 amplicon sequence variants (ASVs) after removal of endosymbiont (*Blattabacterium* sp.) and mitochondrial sequences (Callahan et al., 2016). At the phylum level, at least 80% of each sample was dominated by Bacteroidota, Firmicutes, and Desulfobacterota, in agreement with previous studies on the cockroach gut microbiome (**Figure 1A**) (Tinker and Ottesen, 2016; 2020; Dukes et al., 2023). The relative abundances of these three phyla were similar across all samples excluding xylan-fed cockroaches; these insects hosted notably more Firmicutes and less Desulfobacterota than cockroaches fed other diets (**Figure 1A**).

**Figure 1:**
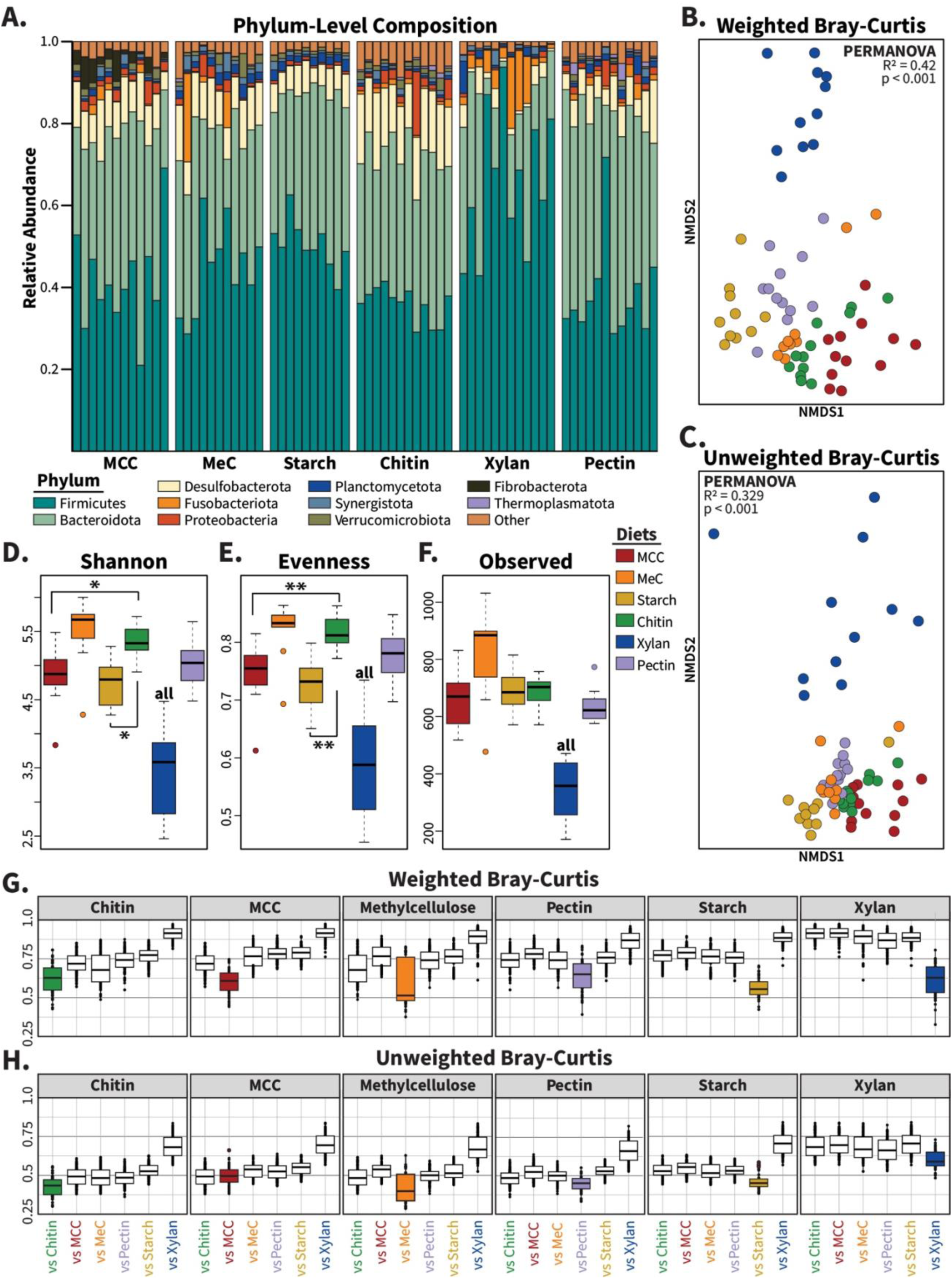
Composition of gut microbiomes from cockroaches fed synthetic diets. (**A**) Barplot showing the relative abundance of phyla across samples for each of the synthetic carbohydrate diets. Bars represent individual hindgut samples, clustered and labeled by diet polysaccharide source. Phyla present at an abundance greater than 1% in at least one sample are plotted. Non-metric multidimensional scaling (NMDS) was used to plot **(B)** weighted and **(C)** unweighted Bray-Curtis dissimilarity. The alpha diversity measures **(D)** Shannon index, **(E)** Pielou’s evenness, and **(F)** number of observed taxa were plotted. Boxplots of **(G)** weighted and **(H)** unweighted Bray-Curtis display each diet vs self (colored boxes) and the other five synthetic diets (white boxes). Samples were rarefied to a constant depth of 7924 sequences for alpha and beta diversity calculations. For alpha diversity measures, pairwise statistics were calculated with Wilcoxon rank-sum tests and multivariate analysis was performed using Kruskal-Wallis tests. PERMANOVA was used to generate statistics for ordination analyses. “all” indicates p<0.05 vs all other diets; * = p<0.05; ** = p<0.01

Alpha diversity, as measured by Shannon index, evenness, and community richness significantly differed across diet treatments (**Figure 1D-F**; Kruskal Wallis p< 0.001 for each). Pairwise analyses found that chitin-fed insects possessed higher Shannon index values (p < 0.05) and community evenness (p < 0.01) than that of MCC- and starch-fed insects, while the xylan diet resulted in lower alpha diversity measures than all other diets (p < 0.05 for each).

Beta diversity analyses using weighted and unweighted Bray Curtis dissimilarity revealed significant impacts of our synthetic diets on gut microbiome composition. On average, between-diet variation was greater than within-diet variation (**Figure 1G and 1H)**, with xylan-fed communities producing distinct shifts from each other diet. Ordination analyses using non-metric multidimensional scaling (NMDS) and PERMANOVA analysis showed that samples clustered based on diet composition in both weighted (**Figure 1B**; PERMANOVA: R^2^=0.42; p<0.001) and unweighted (**Figure 1C**; PERMANOVA: R^2^=0.329; p<0.001) Bray-Curtis metrics, with especially clear separation of the xylan-based diet from other synthetic diets. Removing xylan-fed samples from diversity calculations did not eliminate diet-based clustering for weighted (**Figure S1A;** PERMANOVA: R^2^=0.343; p<0.001) or unweighted (**Figure S1B;** PERMANOVA: R^2^=0.247; p<0.001) measures, suggesting that each carbohydrate source enriched for a unique community composition.

### 4.2 Diet-characteristic taxa enriched by polysaccharide source

We used DESeq2 to identify 76 microbes that exhibited significantly higher abundance in a single synthetic diet across pairwise comparisons against all other treatments, which we termed “diet-characteristic taxa” (**Figure S2**) (Love et al., 2014). Diet-characteristic ASVs were primarily assigned to Firmicutes (n=48) and Bacteroidota (n=20); other phyla with diet-responsive taxa include Fusobacteriota, Deferribacterota, Desulfobacterota, Fibrobacterota, and Spirochaetota. We found that the chitin and methylcellulose diets were not associated with any diet-characteristic taxa by this definition, while diets made with xylan, MCC, starch, and pectin enriched for 45, 10, 13, and 8 ASVs respectively.

### 4.3 Cohort effects on diet-driven differences in gut microbiome composition

To confirm that synthetic diets do produce such dramatic shifts in the community, we evaluated possible cohort effects on gut microbiome responses by replicating and retesting the MCC- and xylan-based diets. These diets were selected for follow-up experiments due to both their contrasting molecular structure and the dissimilarity they generated in Bray-Curtis analyses (**Figure 1B-C**). Data from the first and second experiments exhibited similar alpha diversity measurements (**Figure 2A-C**), with replicates maintaining the significant shifts in alpha diversity (p < 0.001) observed in the initial experiment between these two diets while showing no difference between replicates. Beta diversity analysis showed that samples clustered by both replicate and diet (**Figure 2D-E**). Diet had large effects on both weighted (PERMANOVA: R^2^=0.34; p<0.001) and unweighted (PERMANOVA: R^2^=0.20; p<0.001) Bray-Curtis dissimilarity, while replicate explained minimal effect sizes of 3.7% and 5.8% for weighted and unweighted measures respectively, with only unweighted reaching significance (p < 0.01).

**Figure 2:**
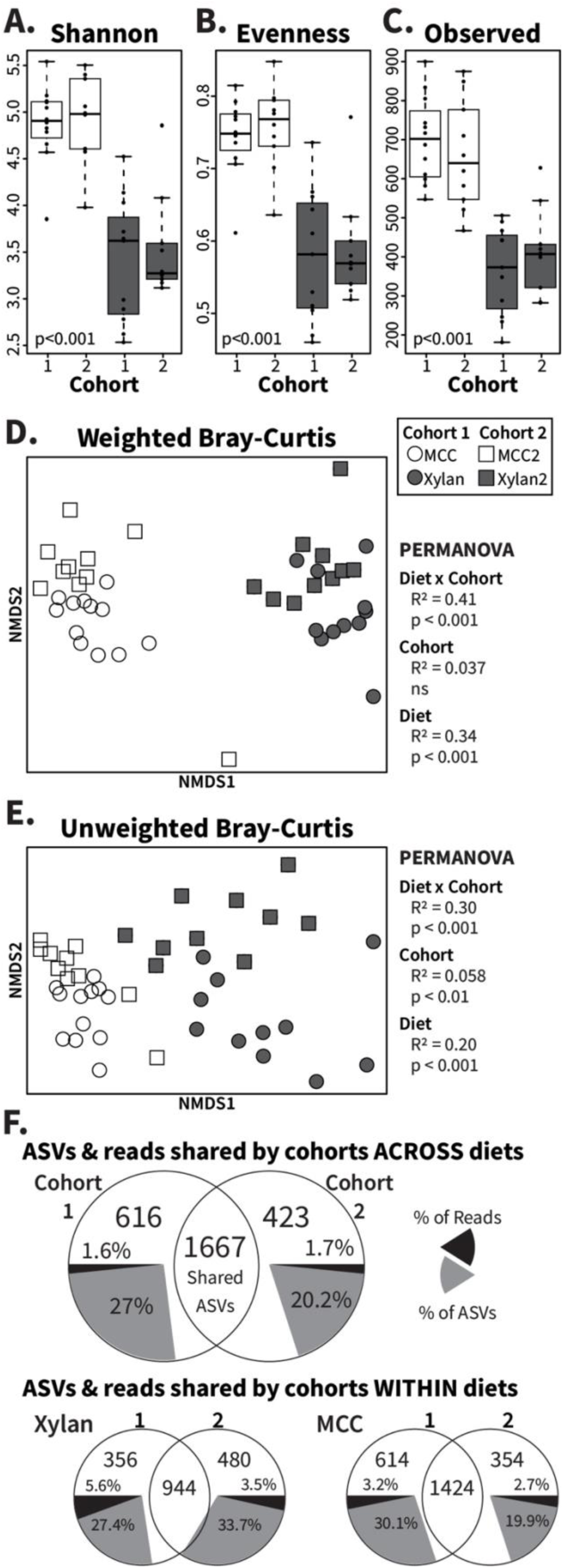
Analysis of cohort effects on gut microbiome responses to MCC and xylan diets. Xylan and MCC-fed samples from replicate experiments were rarefied to 9685 ASVs for alpha and beta diversity assessment. Boxplots show **(A)** Shannon index, **(B)** Pielou’s evenness, and **(C)** number of observed ASVs with Kruskal-Wallis p-values calculated across all individual groups. PERMANOVA was used to calculate R^2^ and p-values for diet (“MCC” and “Xylan”), cohort (“Cohort 1” and “Cohort 2”), and diet x cohort for NMDS ordinations of **(D)** weighted and **(E)** unweighted Bray-Curtis dissimilarity. The last panel **(F)** contains Venn diagrams of shared and unique ASVs between cohorts for both diets together as well as separately, constructed using rarefied count tables. Grey pie slices represent the percent of ASVs observed that are cohort-unique, while black pie slices represent the percentage of sequence reads assigned to the indicated unique ASVs. Ns =no significance

The slight difference observed between the effect of replicate on unweighted vs weighted dissimilarity is reflected by Venn Diagram comparisons (**Figure 2F**) of the ASVs present/absent in the individual cohorts and of the percent of sequences the different sets comprised. While 616 ASVs (27%) were unique to cohort 1 and 423 ASVs (20.2%) unique to cohort 2 (**Figure 2F**, grey pie slices), these ASVs represented only a fraction of the total sequences (**Figure 2F**, black pie slices) obtained from each cohort. Further, 66.7% and 61% of these taxa appeared in only one sample from cohorts 1 and 2 respectively (**Figure S3**), indicating that the majority of differences in composition due to time between studies stem from transient, rare taxa. Separating the diets for these comparisons confirmed the overall findings, with rare taxa contributing few sequencing reads despite comprising 19.7-33.7% of unique ASVs (**Figure 2F**). Altogether, these results show that synthetic diets reproducibly alter the gut microbiome composition in cockroaches.

### 4.4 Testing the impact of alternative diet formulations

To confirm the role of carbohydrates as key modulators of the gut community in our synthetic diets, we used xylan and MCC-based diets to test the impact of removing protein or micronutrients, and the replacement of long-chain fiber with component sugars influenced the gut microbiome. For protein-deficient diets (**Table 1**), casein and peptone were replaced by mass with either xylan or MCC, while in vitamin-deficient diets, both the vitamin mixture and cholesterol were replaced with additional carbohydrate. Three simple sugar diets featuring mono/disaccharides derived from xylan and MCC were also tested: xylose, cellobiose, and glucose.

Weighted Bray-Curtis ordination analysis of cockroaches fed these diets and standard xylan/MCC synthetic diets (**Figure 3A**) revealed that both xylan-fed and MCC-fed samples clustered by polysaccharide regardless of vitamin or protein content and separately from sugar diets (PERMANOVA: R^2^=0.434; p<0.001). In unweighted analyses (**Figure 3B),** MCC-associated clusters overlapped with sugar diets, but retained separation from xylan-fed samples (PERMANOVA: R^2^=0.36; p<0.001). The alpha diversity profiles of the deficient diets matched the standard MCC or xylan diets as well, as measured by Shannon index (**Figure 3C**), evenness (**Figure S4A**), and number of observed ASVs (**Figure S4B**). The cellobiose communities displayed slightly higher Shannon index values than the standard and protein-deficient MCC communities, while xylose-fed insects possessed noticeably more even and diverse communities than the xylan diets. Likewise, community composition at the genus level (**Figure 3D**) showed little difference between insects fed diets sharing polysaccharide source with or without all ingredients, with sugar-fed microbiota reflecting each other more than the carbohydrate they are derived from. At the ASV level (**Figure S5**), xylose was found to enrich for an abundant *Lachnoclostridium* ASV that is heavily associated with xylan, unlike cellobiose or glucose, which showed little representation of MCC-associated taxa such as *Fibrobacter* and *Ruminococcus*.

**Figure 3:**
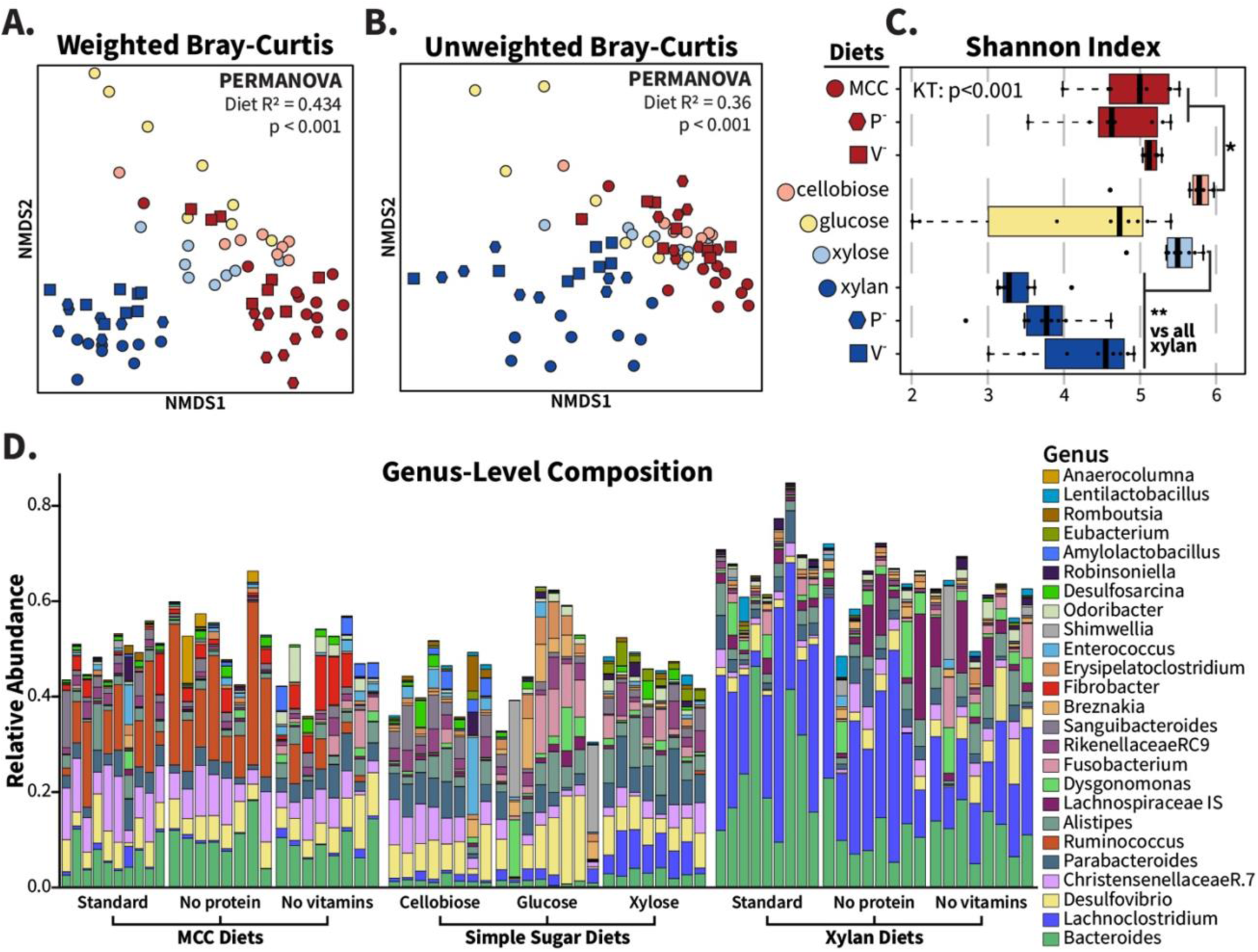
Fiber source, not protein, vitamins, or sugar composition, determines community structure from xylan and MCC-based synthetic diets. Deficient and simple sugar variations of MCC and xylan synthetic diets were fed to adult cockroaches for two weeks, and hindgut community compositions were compared with replicated xylan and MCC samples. For these analyses, samples were rarefied to 12274 reads. NMDS ordinations were made for **(A)** weighted and **(B)** unweighted Bray-Curtis dissimilarity, and PERMANOVA used to calculate R^2^ and p-values with “diet” as the grouping factor. Alpha diversity is displayed via **(C)** Shannon index with Wilcoxon rank-sum test used for pairwise comparisons. The relative abundance of abundant genera found in the MCC- and xylan-based diets are visualized in **(D).** * =p <0.05; ** =p<0.01.

### 4.5 Comparison with whole food diets

The changes in community composition triggered by our synthetic diets were unexpected given that previous experiments examining the impact of whole food diets with strongly differing macronutrient profiles did not induce substantial change in gut microbiome composition (Tinker and Ottesen, 2016). Therefore, we compared the samples from this current study (“synthetic” diet type) to samples from the previous study (“whole food” diet type) of cockroaches fed butter, tuna, honey, white flour, or whole wheat flour (Tinker and Ottesen, 2016).

We found that gut microbiome samples from cockroaches fed synthetic diets exhibited higher ASV richness (p<0.01) but lower evenness (p<0.001) and Shannon index (p<0.01) than those from insects fed whole foods (**Figure S6A-C**). Synthetic and whole food diets produced distinct diet type clusters in NMDS ordinations (**Figure 4A-B**) for weighted (PERMANOVA: R^2^=0.105; p<0.001) and unweighted (PERMANOVA: R^2^=0.152; p<0.001) Bray-Curtis dissimilarities. When we analyzed the samples by diet, we found diet explained more variation in NMDS ordination than diet type as interpreted from PERMANOVA R^2^ values (weighted: R^2^=0.393; unweighted: R^2^=0.369).

**Figure 4:**
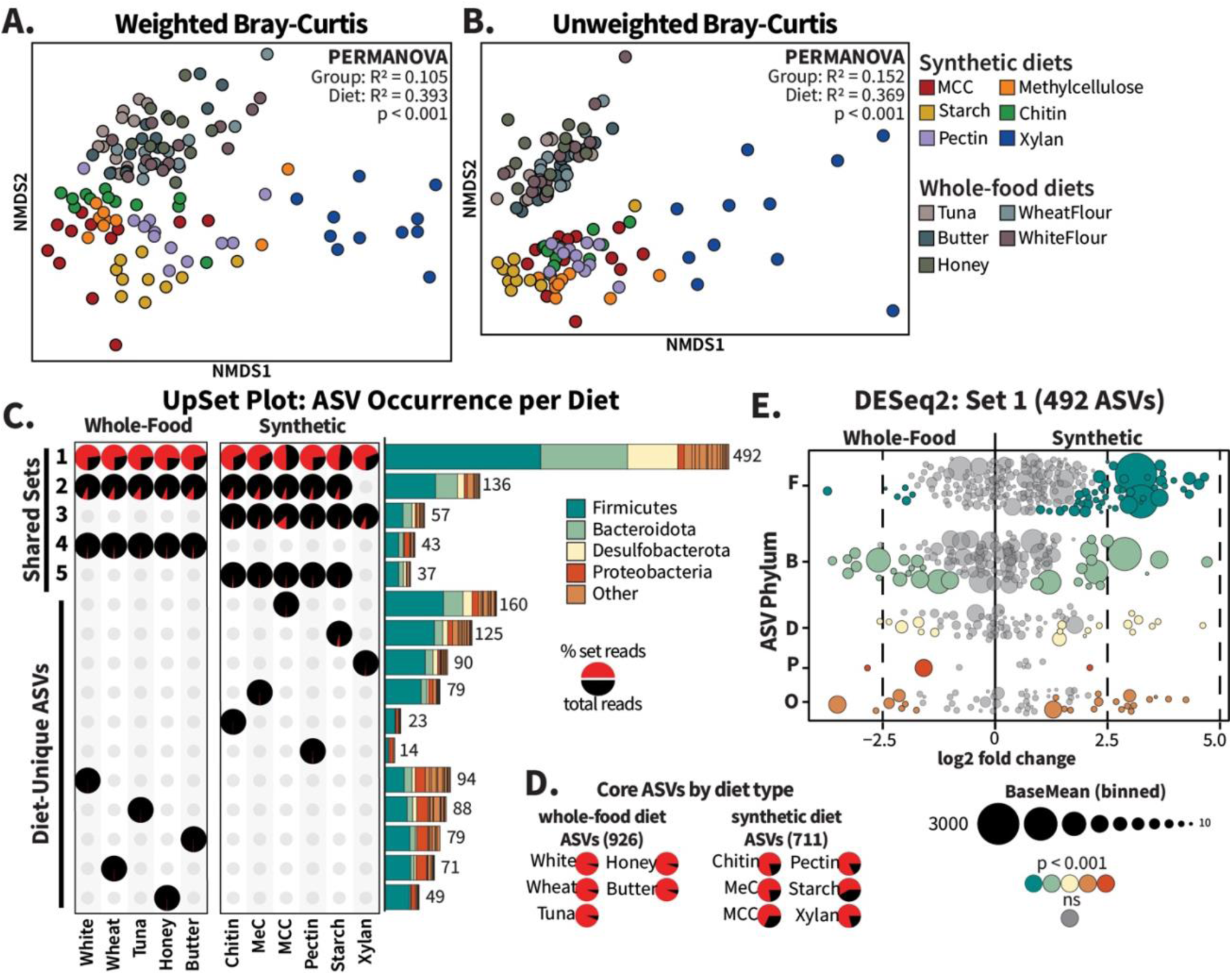
Whole-food diets share more ASVs than synthetic diets. Raw sequence data from Tinker and Ottesen (2016) for cockroaches fed tuna, butter, honey, wheat flour, and white flour (“Whole Food Diets”) were reprocessed using the methods in this experiment to generate comparable ASVs. For beta diversity comparison, all samples were rarefied to 7924 ASVs and NMDS ordinations were generated for **(A)** weighted and **(B)** unweighted Bray-Curtis distances; R^2^ and p-values for diet type comparisons were calculated using PERMANOVA. For **(C)** UpSet plot analysis, the five largest intersections (Sets 1-5) and diet-unique sets are displayed. Pie charts represent the percent of reads within a diet that originate from each set (red slices), and the bar charts are colored to display the phylum-level distribution of ASVs assigned to each set. Read abundance of core ASVs, or those present in all synthetic or whole food diets regardless of presence in the other diet type, are visualized in **(D)** pie charts per diet. For the MA plot in panel **(E)**, raw sequence count tables for the ASVs identified as “Set 1” were analyzed using DESeq2 with diet type as the design factor. The ASV circles are scaled according to baseMean size and colored by phylum.

Beta dispersion analysis of variation within diet types showed that, together, the gut microbiota of cockroaches fed synthetic diets was more variable than that observed among whole food fed cockroaches (**Figure S7A-B**; Tukey’s HSD: p<0.001). However, when diets were analyzed individually, they were equally dispersed in weighted Bray-Curtis dissimilarity (**Figure S7A)** but not unweighted measures **(Figure S7B;** ANOVA: p<0.001). Xylan-fed cockroaches exhibited significantly greater within-group variability than all ten other diets (Tukey’s HSD range: p = 0.037 – 3.17e^-06^), with no significant differences observed in pairwise comparisons of all other diets. When synthetic diets were compared to whole food diets without including xylan-fed samples, we observed no significant differences in unweighted beta dispersion (**Figure S7D**) or Shannon index values (**Figure S6B, red boxes**). However, significant differences remained in weighted Bray-Curtis dispersion (**Figure S7C**), richness and evenness (**Figure S6A, S6C**), and in both weighted and unweighted Bray-Curtis ordination analysis **(Figure S8A-B)**, highlighting that the altered microbiomes produced by the synthetic diet type were not solely due to biases produced by xylan-fed samples.

To verify that inclusion of whole food dietary components alone was not sufficient to eliminate fiber-dependent gut microbiome shifts, we tested the impact of diets mimicking our synthetic diets but with the purified protein components replaced with canned tuna. These diets induced community composition shifts similar to those observed in carbohydrate matched diets containing purified proteins rather than supporting protein-associated communities (**Figure S9**). Xylan-containing diets generally produced communities with lower alpha diversity scores (**Figure S9D-F**) and clustered away from MCC-containing diets and dog chow-fed insects we included as controls in weighted (**Figure S9B;** PERMANOVA: R^2^=0.395; p<0.001) and unweighted (**Figure S9C;** PERMANOVA: R^2^=0.392; p<0.001) analyses. Despite the discordant structural complexities between tuna fish and purified casein/peptone amino acids, the protein portion of the synthetic diets exerted less influence than carbohydrate source.

### 4.6 Core taxa differences between synthetic and whole food diets

Given these strong differences in community structure, we utilized the R package UpSetR to determine how the ASVs in different diet types overlap (Conway et al., 2017). UpSet plots are akin to Venn diagrams, considering only presence/absence of an ASV. Rarefied count tables were aggregated by diet and ASVs were marked as either present or absent per diet. ASVs present in the same subset of diets were grouped into “Sets”, with the phylum-level composition per set depicted as stacked bar charts labelled with the number of included ASVs (**Figure 4C**). We supplemented the UpSet plot with pie charts illustrating the relative abundance (calculated as the fraction of total reads recovered from the collapsed treatment group) of reads assigned to ASVs within each set (**Figure 4A**), in addition to pie charts representing “core” ASVs present in all whole-food or all synthetic diets, regardless of presence in the other dietary group (**Figure 4D**). For simplicity’s sake, **Figure 4** shows only the five largest intersecting sets as well as all single-diet sets; additional sets are presented in **Figure S10**.

A total of 492 ASVs (“Set 1”) were shared across all diet treatments (**Figure 4C**). These ASVs made up over half of the sequences recovered for all diets except the MCC diet, for which they represented 49% of sequences (**Figure 4C**, pie charts). Only 43 ASVs (“Set 4”) were exclusive to the whole food diets, contributing between 0.9% and 1.65% of reads in these diet sets. The 57 ASVs (“Set 3”) identified as exclusive to synthetic diets made up 1.6-3.4% of the reads recovered from starch-, pectin-, chitin-, and methylcellulose-fed cockroaches, and 7% of xylan- and 13% of MCC-fed cockroaches. Together, these results argue that the synthetic diets did not eliminate core taxa present in the guts of cockroaches fed whole foods, nor did they result in hindgut colonization by a large new set of microbial taxa.

Similarly, individual synthetic diets were not associated with hindgut colonization by large groups of unique microbes. In general, taxa that were unique to individual synthetic or whole food diets represented a very small proportion of sequences recovered (**Figure 4C**). ASVs unique to the MCC diet formed the second largest set overall, with 160 diet-unique taxa, yet they only represented 0.4% of total recovered sequences (**Figure 4C**). A xylan-based diet, which repeatedly produced the largest community dissimilarities (**Figure 4A-B; Figure S7**), was associated with 90 diet-specific taxa comprising only 0.63% of reads. Interestingly, our analysis revealed that a xylan-based diet did result in the loss of 136 taxa that were present in all other diets in abundances ranging from 4.78%-10.14% (“Set 2” in **Figure 4C**). However, other sets that excluded individual diets were substantially smaller (**Figure S10A**) suggesting that this was not a common mechanism underlying the diet-driven changes in gut microbiome composition. Instead, synthetic diet-driven changes in gut microbiome composition were primarily associated with changes in the relative abundance of individual taxa that were consistently present in the cockroach gut regardless of dietary treatment.

To follow up on these observations, we used DESeq2 to assess enrichment of individual ASVs between synthetic vs whole food fed cockroaches. Of the 492 ASVs present in at least one sample belonging to each diet (Set 1), 95 ASVs were significantly enriched in cockroaches fed synthetic diets, while 38 ASVs were enriched in cockroaches fed whole food diets (**Figure 4E**; padj < 0.001). Set 1 ASVs were present in all diets, often at high abundances, and shifts across diet type were modest (log fold change <5). Bacteroidota, Desulfobacterota, Proteobacteria and other phyla showed similar enrichment distributions to each other, but more Firmicutes were enriched by the synthetic diets.

We also examined enrichment of ASVs that fell outside of Set 1 but were both abundant (baseMean > 1) and present in at least 5 samples (**Figure S11**). These 1270 ASVs generally had smaller abundances than members of Set 1 (**Figure 4E**), but greater log fold changes between diet types. The ASVs *Ruminococcus_NA* and *Fusobacterium ulcerans* were exceptions, being both highly abundant (baseMean = 1020 and 651, respectively) and unique to the synthetic diets. In contrast, the *Christensenellaceae* ASV (R7_NA.68) unique to whole food diets had high log-fold change, but a baseMean of only 4.29. Apart from a few highly abundant ASVs in the synthetic diets, most abundant, diet-enriched taxa were shared by all diets, supporting that change is driven by common gut bacteria restructuring the community rather than interloping bacteria disrupting the community.

### 4.7 Differential diet-based fluctuations of abundant Firmicutes and Bacteroidota ASVs

To explore the impact of diet on microbial taxa associated with fiber degradation, we evaluated dietary responses of abundant taxa within the Firmicutes and Bacteroidota. For this analysis, we selected the two most abundant representatives of each of these phyla from each diet and examined their abundance across all diets (**Figure 5**). We identified 16 Firmicutes and 11 Bacteroidota as most or second-most abundant in at least one diet.

**Figure 5.**
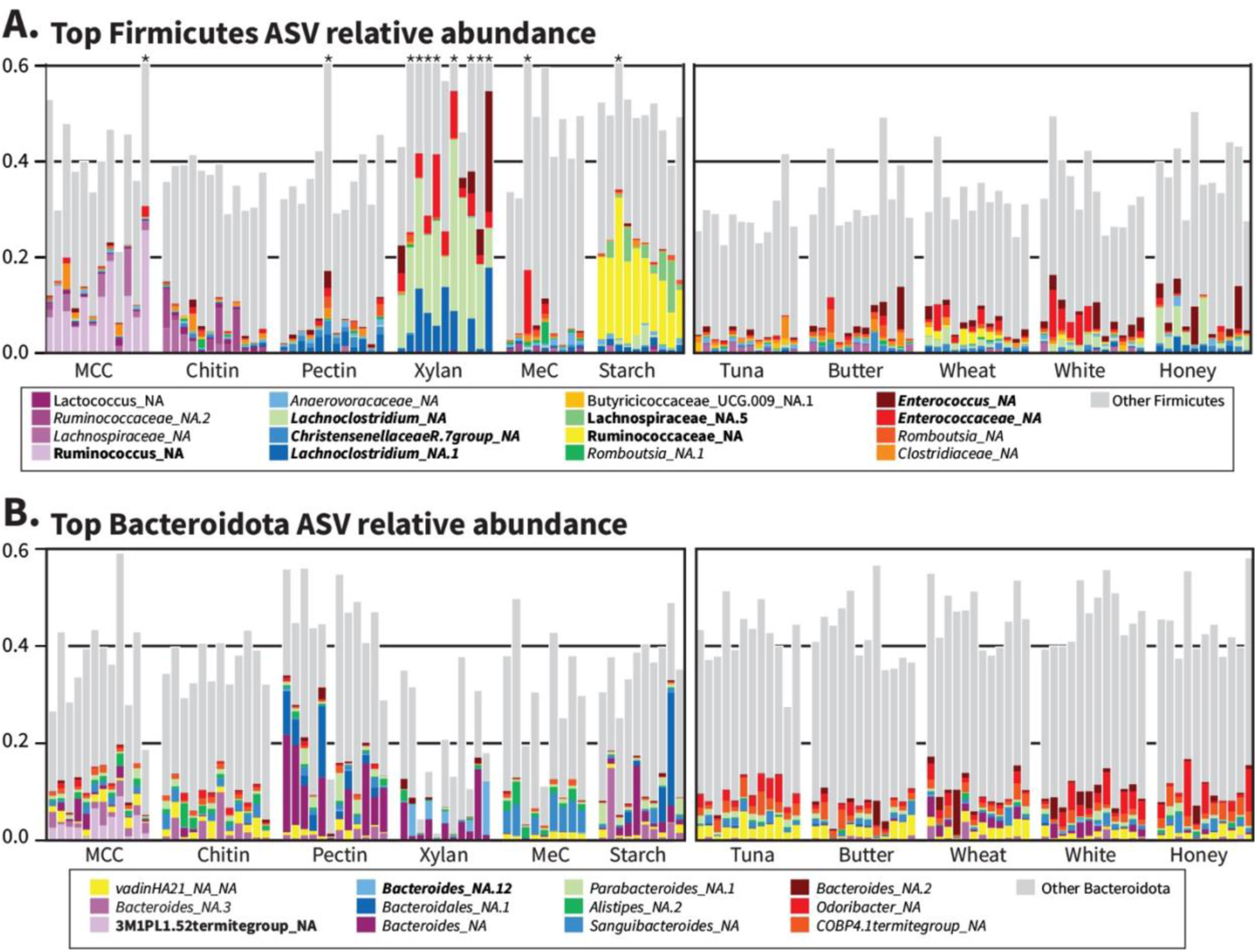
Individual ASVs explain large differences between synthetic but not whole-food diets. Variance-stabilized count data from DESeq2 were used to determine the top two ASVs for every diet belonging to **(A)** Firmicutes and **(B)** Bacteroidota, and the relative abundances of ASVs belonging to the combined ‘top ASV’ set were plotted for all samples. Grey bars include all Firmicutes or Bacteroidota not named in the key. * indicates “Other Firmicutes” extends beyond 60% relative abundance; please refer to Figure 1A for full values. Bolded names indicate diet-characteristic ASVs from **Figure S2**, and italics indicate Set 1 ASVs from Figure 4C.

The most abundant taxa from both Firmicutes and Bacteroidota represented a small fraction of reads across the whole foods diets, consistent with the higher Shannon diversity and evenness of the gut microbiome for whole food-fed cockroaches (**Figure S6B, C**). In contrast, the synthetic diets produced strong ‘blooms’ of individual ASVs, particularly among the Firmicutes (**Figure 5A**).

Several abundant Firmicutes were both present in Set 1 (**Figure 4C**) and enriched in individual synthetic diets (**Figure S2**), namely *Lachnoclostridium_NA* (xylan), *Lachnoclostridium_NA.1* (xylan), *Enterococcus_NA* (xylan), *Enterococcaceae_NA* (xylan), and *ChristensenellaceaeR7_NA* (pectin). In contrast, Firmicutes identified as enriched in the MCC and Starch diets were not typically found across all diets: *Ruminococcus_NA* (MCC), *Lachnospiraceae_NA.5* (starch), and *Ruminococcaceae_NA* (starch). Among the Bacteroidota (**Figure 5B**), we observed greater overlap in the most abundant taxa present in each diet group, with all but one (*3M1PL1.52termite_NA*) of the abundant Bacteroidota ASVs belonging to Set 1, and only two classified as diet-characteristic: *3M1PL1.52termite_NA* (MCC) and *Bacteroides_NA.12* (pectin).

### 4.8 Analysis of Microbial Co-Correlation Networks

We constructed co-correlation networks with SparCC to examine the community structure underlying synthetic and whole food microbiome data sets (Friedman and Alm, 2012). To filter out noise and reduce spurious correlations, datasets were filtered to include ASVs present in at least 25% of samples for each diet type, resulting in 976 ASVs for whole food diets and 700 ASVs for synthetic diets. Networks were further pruned to contain edge weights with absolute values larger than 0.4, removing 75 and 168 ASVs from whole food and synthetic networks respectively. Both positive and negative edges were retained for network layout formation (**Figure S12**), but for analysis, only positive edges were considered (**Figure 6**). After negative edge removal, the whole food network contained 875 ASVs with 9515 edges forming two connected components (**Figure 6A**), while the synthetic diet network contained 497 ASVs with 2536 edges that formed six connected components (**Figure 6B**). Networks at SparCC correlation levels of 0.5, 0.6, and 0.7 are included in **Figure S13**.

**Figure 6:**
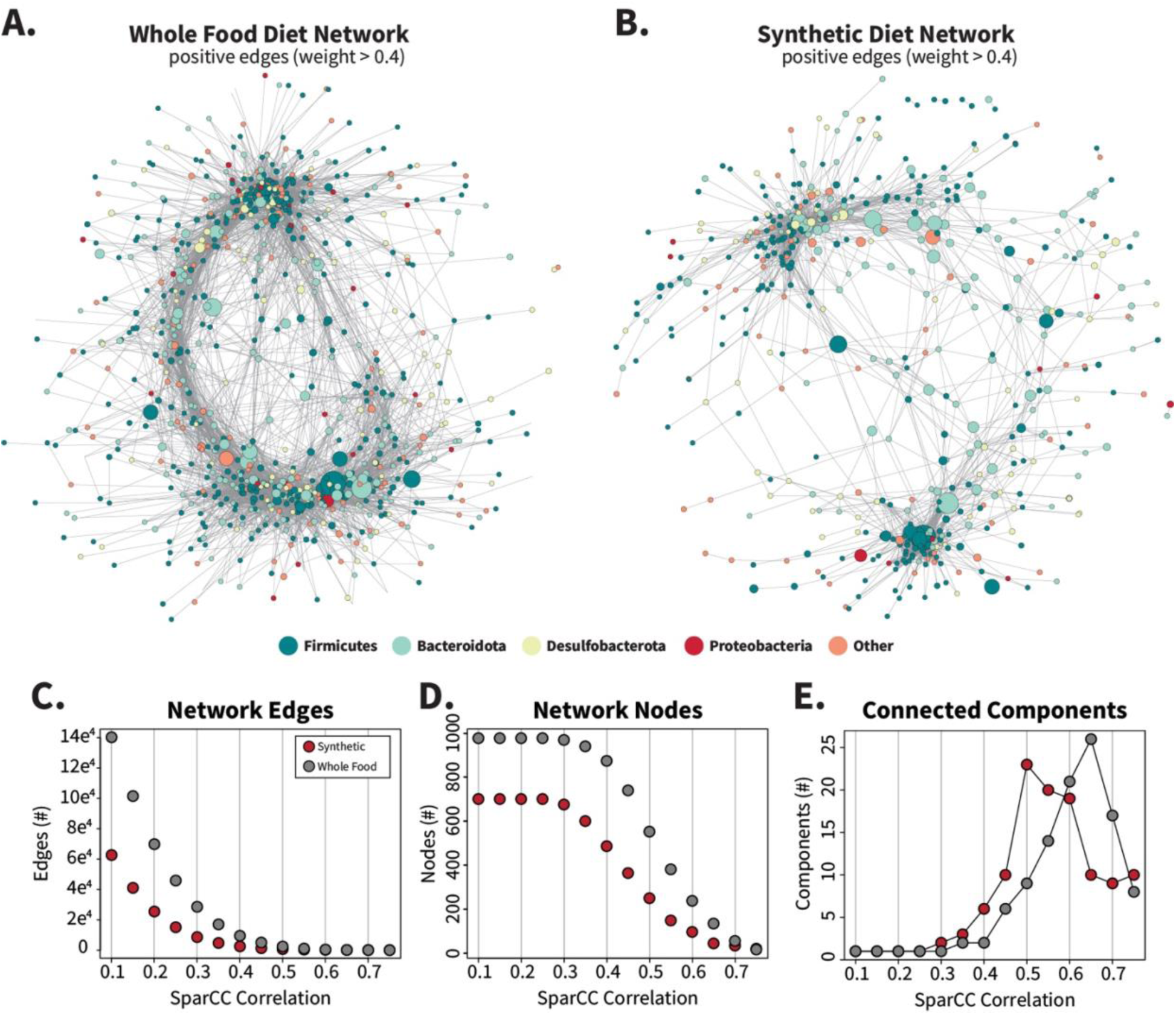
Synthetic diet correlation networks are smaller and less interconnected than whole food diet correlations networks. Networks were calculated by SparCC from filtered count tables (ASVs present in > 25% of samples per diet set) for synthetic and whole food diets separately to create two distinct networks containing 976 (whole food) or 700 (synthetic) nodes. Networks were imported into Cytoscape and edges with absolute values < 0.4 were removed to generate panels **(A)** and **(B)**. Negative edges were included during initial layout generation with the edge-weighted spring embedded layout method and are displayed in **Figure S12.** The number of nodes, edges, and connected components that remain when the networks are filtered by increasing correlation values are charted in **C**, **D,** and **E** respectively.

Gut microbiota from whole food-fed cockroaches formed an extensive and dense interaction network (**Figure 6A**), with higher edge counts (**Figure 6C**), node counts (**Figure 6D**), node degree (**Figure S14A**) network strength (**Figure S14B**), and betweenness scores (**Figure S14C-D**) than synthetic diets at most levels of filtering based off SparCC correlation values. High-degree nodes, or ASVs with large numbers of neighbors, were present throughout the whole-food network structure, while in the synthetic diets two primary clusters of ASVs appeared with fewer connecting ASVS. When a range of inclusion cutoffs were considered, the synthetic diet network degraded more quickly than the whole food network into separate connected components (**Figure 6E, Figure S13)** displaying greater fragility and tendency to fragment. These overall network structures suggest that the synthetic diets disrupt the stability of the cockroach microbiome.

## 5 Discussion

The core finding of this work is that synthetic diets dominated by a single carbohydrate type induced marked fiber-dependent changes in hindgut microbial community composition in omnivorous cockroaches. Diets featuring the hemicellulose xylan induced the largest shifts in alpha diversity, inter-individual variability, and overall community composition (**Figure 1**). Xylan is a common structural polysaccharide containing a xylose backbone with branching arabinose, glucuronic acid, and/or acetyl groups that has been extensively investigated for prebiotic potential both in livestock and humans (Broekaert et al., 2011; Jefferson and Adolphus, 2019; Jana et al., 2021; Smith and Melrose, 2022). Research investigating xylan degradation ability in gut microbiota mainly focuses on *Bacteroides* (Dodd et al., 2011; Zhang et al., 2014; Centanni et al., 2017; Déjean et al., 2019; Kmezik et al., 2021; Pereira et al., 2021) although select Firmicutes such as *Roseburia intestinalis* have been identified as key butyrate-producing xylan fermenters (Mohand-Oussaid et al., 1999; Leth et al., 2018; Hershko Rimon et al., 2021). Interestingly, we found that in cockroaches, xylan-based diets decreased the relative abundance of Bacteroidota, while increasing Firmicutes (**Figure 1A**). Concurrent with this enrichment of specific taxa, xylan decreased the overall diversity of the gut community (**Figure 1**). This result contrasts with studies of gut microbiome composition changes in chickens (Zhou et al., 2021) and mice (Berger et al., 2021) fed diets supplemented with xylooligosaccharides where gut diversity increased, although both studies reflected the increase in *Lachnospiraceae* observed in our experiments.

However, while xylan-based diets induced the largest change in hindgut community composition, samples clustered by diet treatment even when the xylan treatment group was excluded (**Supplement 1**). Most of these diets were associated with ‘blooms’ of specific Bacteroidetes and Firmicutes, excluding those made with methylcellulose and chitin, which were not associated with enrichment of any single taxa in pairwise comparisons with the other diets. This suggests that while the disproportionate impact on community structure due to xylan warrants its own research, synthetic diets in general appear to reduce hindgut microbiome stability compared to whole food-based diets.

These results stand in stark contrast to previous work from multiple investigators, who found minimal to no change in hindgut microbial community composition in response to diet alterations (Schauer et al., 2014; Tinker and Ottesen, 2016; Lampert et al., 2019). A commonality between these experiments is that the investigators utilized complex, minimally processed or whole food diets that were highly biased in macromolecular composition. On the other hand, a few investigators have previously observed substantial influence of diet on the gut microbiome composition (Bertino-Grimaldi et al., 2013; Pérez-Cobas et al., 2015; Zhu et al., 2022). These experiments all utilized synthetic diets that contained purified carbohydrate and protein sources. For example, in experiments using *B. germanica*, Pérez-Cobas et al (Pérez-Cobas et al., 2015) prepared synthetic diets with a dextrin and micronutrient base amended with either 50% cellulose or 50% casein while Zhu et al (Zhu et al., 2022) used diets composed of a cellulose and micronutrient base with supplemented with 40% by mass purified starch, casein, or sesame oil. In *P. americana*, Bertino-Grimaldi et al (Bertino-Grimaldi et al., 2013) utilized purified cellulose to compare with sugarcane bagasse, a complex dietary substrate.

A disproportionate influence of synthetic diets with highly purified components on gut microbial composition has been observed beyond cockroach research as well. Termites, a close relative to cockroaches, responded to single carbohydrate source diets with larger alterations in gut community composition than termites fed mixed-carbohydrate diets (Miyata et al., 2007). Other insect model systems produced similar results, such as in silkworms (Dong et al., 2018), ladybugs (Xie et al., 2024), waxworms (Gohl et al., 2022), and honeybees (Powell et al., 2023). Among mammals, dogs provided a purified diet also exhibited reduced alpha diversity compared to those fed a complex diet (Allaway et al., 2020), wild-caught mice transitioned from natural diets to laboratory diets lost large portions of native gut microbes (Martínez-Mota et al., 2020), and humans given meal replacement shakes showed loss of biodiversity in their microbiome compositions (Gurry et al., 2018).

Comparison of our dataset with data recovered in previous whole food-based dietary experiments suggest that synthetic diets altered the gut community by inducing overgrowth of microbes already present in the cockroach gut microbiome (**Figure 4).** Taxa that were unique to individual diets represented <1% of sequence reads in all diets but starch, of which they made up 2.84% (**Figure 4**). In contrast, 15 out of 20 highly abundant Firmicutes and Bacteroidota associated with one or more diets were shared across all diet types, while the remaining 5 were found sporadically in other diets (**Figure 5)**.

We hypothesize that highly purified synthetic diets enabled microbial ‘specialists’ to bloom beyond their former constraints in the whole food diets. While the whole food diets used in Tinker and Ottesen (Tinker and Ottesen, 2016) were highly biased in macromolecular composition, as natural foods they may have had a more diverse nutritional profile according to the “eye” of a bacterium (Hernot et al., 2008; Centanni et al., 2017; Puhlmann and de Vos, 2022). Both whole and white wheat flour, for example, are composed predominantly of endosperm-derived starch, which physically impacts microbial adhesion due to variance in particle size following milling (Fernando et al., 2012; Lin et al., 2019). The bran, germ, and, to a lesser degree, endosperm fractions of wheat contain structural polysaccharides and bioactive phytochemicals that are targeted by gut bacteria and influence health parameters of the host (Adom et al., 2005; Okarter and Liu, 2010; Cardona et al., 2013; Parkar et al., 2013), while honey contains complex mixtures of sugars (glucose, fructose, disaccharides) and fructooligosaccharides in addition to organic acids, nitrogenous compounds, vitamins, and bee-derived enzymes (Sultana et al., 2022). Tuna and butter are similarly high-complexity substrates that offer resident gut microbes diverse metabolizable compounds that are lost in purified dietary components such as those utilized in our study. The purified components used in our synthetic diets are not entirely homogeneous but are far more processed than the ‘natural’ whole foods used in our comparison sets and lack bioactive compounds present basally in these diets (Puhlmann and de Vos, 2022). As a result, the less complex dietary structures of our synthetic diets may not require as much cooperative metabolism to digest while the overall loss in nutritional diversity edges out bacteria that are unable to keep pace with specialist polysaccharide fermenters.

This conclusion is supported by network analysis of the diet types (**Figure 6**), where the whole food network is highly interconnected with numerous paths from one ASV to the next, while the synthetic diet network is easily fragmented into modules of microbes that are weakly or negatively associated with the other network members (**Figure S13, Figure S14**). Hallmarks of a stable and resilient gut microbiome often include metabolic redundancy, where many microbes are capable of filling different niches and are rate limited by their partner activity but can adjust to shifts in diet without extreme disruption (Ze et al., 2013; Pereira and Berry, 2017). Synthetic diets may weaken the cooperative potential of the gut microbiome, narrowing the niches that can flourish to fewer bacteria, and allow drastic shifts such as those observed in the xylan diet that would otherwise not occur.

A key limitation of this study is the fact that the comparison group of “whole food” fed cockroaches were from an earlier experiment and we lack ‘contemporaneous’ controls fed whole food diets. However, an examination of cohort effects suggest that observed responses to synthetic diets were highly conserved across cohorts in experiments conducted one year apart (**Figure 2**). Further, despite the large time gap between the whole food and synthetic diet experiments, a set of 492 ASVs shared by all treatment groups comprised approximately 68% of the reads recovered (**Figure 4**), suggesting minimal drift in overall gut microbiome composition due to time alone.

Overall, this study showed that synthetic diets that were highly enriched in a single carbohydrate type could induce large shifts in gut microbiome composition in the American cockroach, which has previously been shown to be highly resistant to diet-induced shifts in gut microbiome composition (Tinker and Ottesen, 2016). Changes in gut microbiome composition in response to synthetic diets were primarily associated with ‘blooms’ of diet responsive ASVs already present at lower levels in the gut rather than introduction new taxa. This alteration in gut microbiome composition is associated with fragmentation of gut microbiome interaction networks and increased inter-individual variability. Together, these results suggest that overconsumption of a single, purified class of polysaccharides can have destabilizing effects on the gut microbiota. This has major implications for the potential for prebiotics to induce changes in gut microbiome composition even in highly stable systems, as well as potential implications regarding the impact of highly processed foods on gut microbiome homeostasis. Future work will explore the functional and metabolic basis of observed shifts in microbial community composition.

## Supporting information

Compiled Supplemental Figures and Tables

## 6 Data Availability

Data associated with this study are available from the NCBI short-read database under BioProjects PRJNA1096047 and PRJNA1105088. Data associated with the earlier study of the impact of whole food diets on cockroaches is available under BioProject accession PRJNA320546.

## 8 Acknowledgements

We thank Darian Talamantes for assistance with initial diet formulation, and Dr. Kara Tinker for performing the foundational work serving as the basis of this experiment. We also thank Sarah Beth Griffin for her assistance with cockroach husbandry, dissections, and preparing samples for DNA sequencing. A previous version of this manuscript was submitted as a preprint under the title “Synthetic diets containing a single polysaccharide disrupt gut microbial community structure and microbial interaction networks in the American cockroach” (Dockman and Ottesen, 2024).

