## Supplementary material for "Purified fibers in chemically defined synthetic diets destabilize the gut microbiome of an omnivorous insect model": Compiled Supplemental Figures and Tables

#### Supp Fig 1

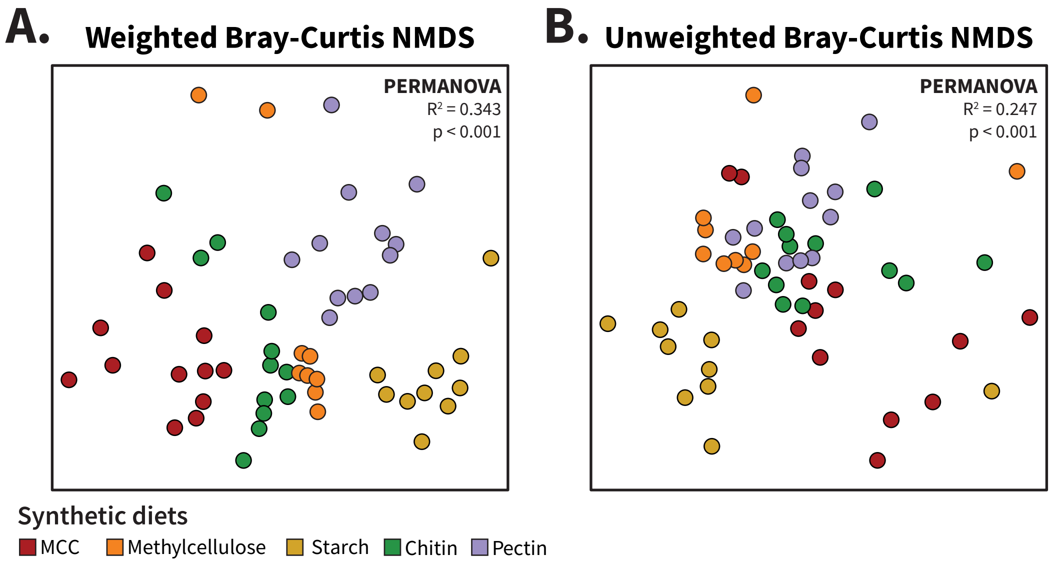


**Supplement 1: NMDS ordination analysis of synthetic diet samples excluding xylan-fed cockroaches.** As in Figure 1, samples were rarefied a constant depth of 7924 sequences for alpha and beta diversity calculations. Non-metric multidimensional scaling (NMDS) was used to plot **(A)** weighted and **(B)** unweighted Bray-Curtis dissimilarity of gut communities from synthetic diets, excluding those fed the xylan diet. Multivariate analysis was performed using PERMANOVA.

### Supp Fig 2

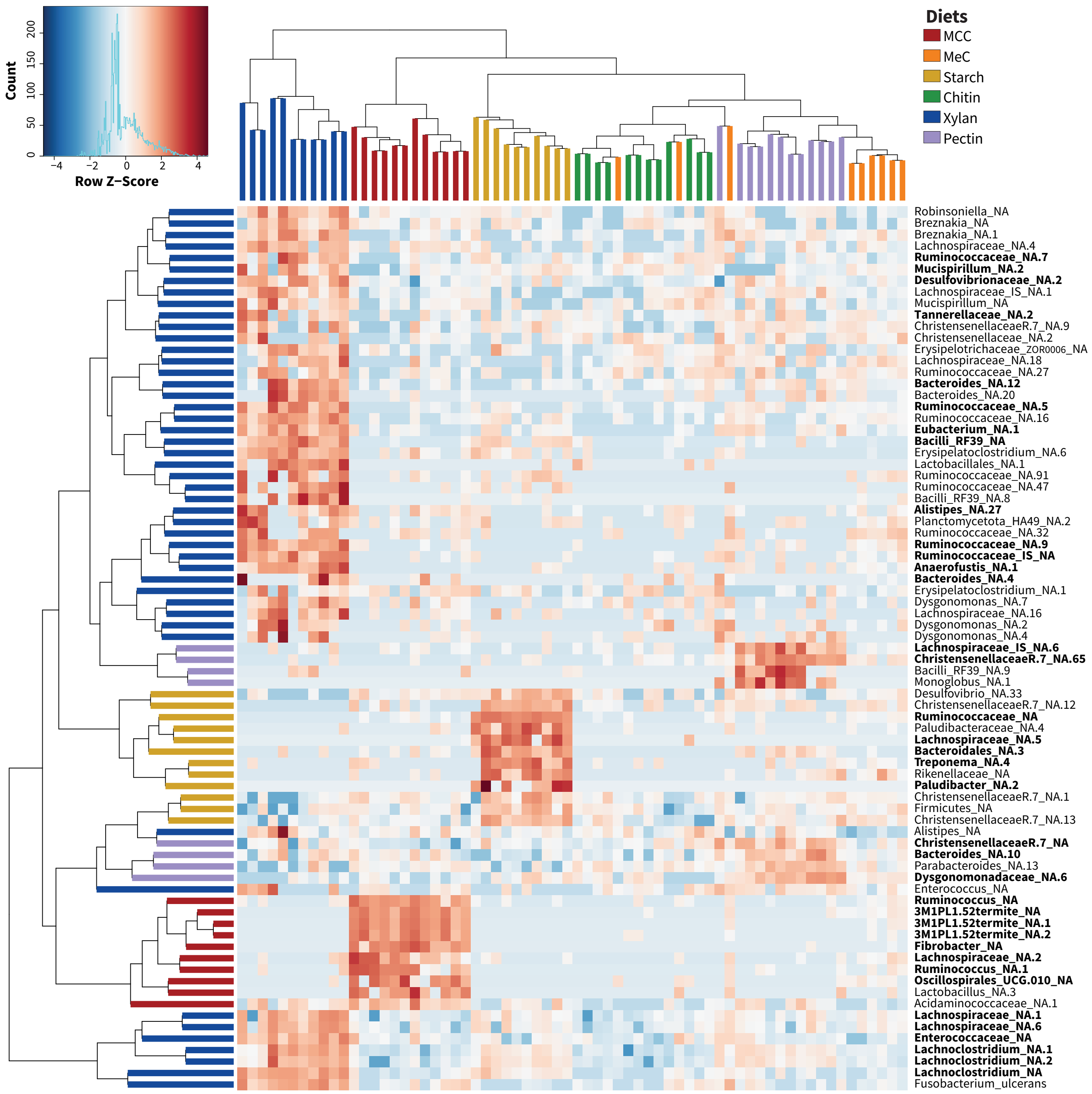


**Supplement 2: Diet-characteristic ASVs determined by DESeq2.** Pairwise DESeq2 analysis was performed on filtered (present in 5 or more samples) raw counts for all synthetic diets (n=66), and ASVs significant in one diet vs all others at adjusted $p < 0.05$ with a baseMean higher than 10 were selected (76 total). Variance stabilized transformed counts obtained from DESeq2 were scaled by row to generate z-scores for plotting. Dendrograms are colored by diet per sample (column) and diet the ASVs are associated with (row). Bolded names indicate the ASV belongs to "Set 1" as presented in Figure 4.

### Supp Fig 3

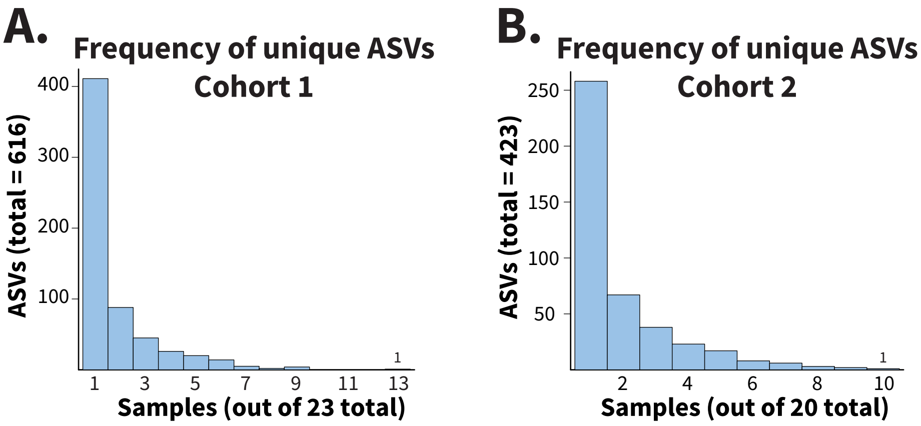


**Supplement 3 : Most unique ASVs in the Xylan/MCC replicate experiments are singletons.** Unique ASVs from the rarefied **(A)** Cohort 1 and **(B)** Cohort 2 data were assessed for frequency of occurrence using the histogram function in R. Samples were rarefied to 9685 ASVs for comparison.

#### Supp Fig 4

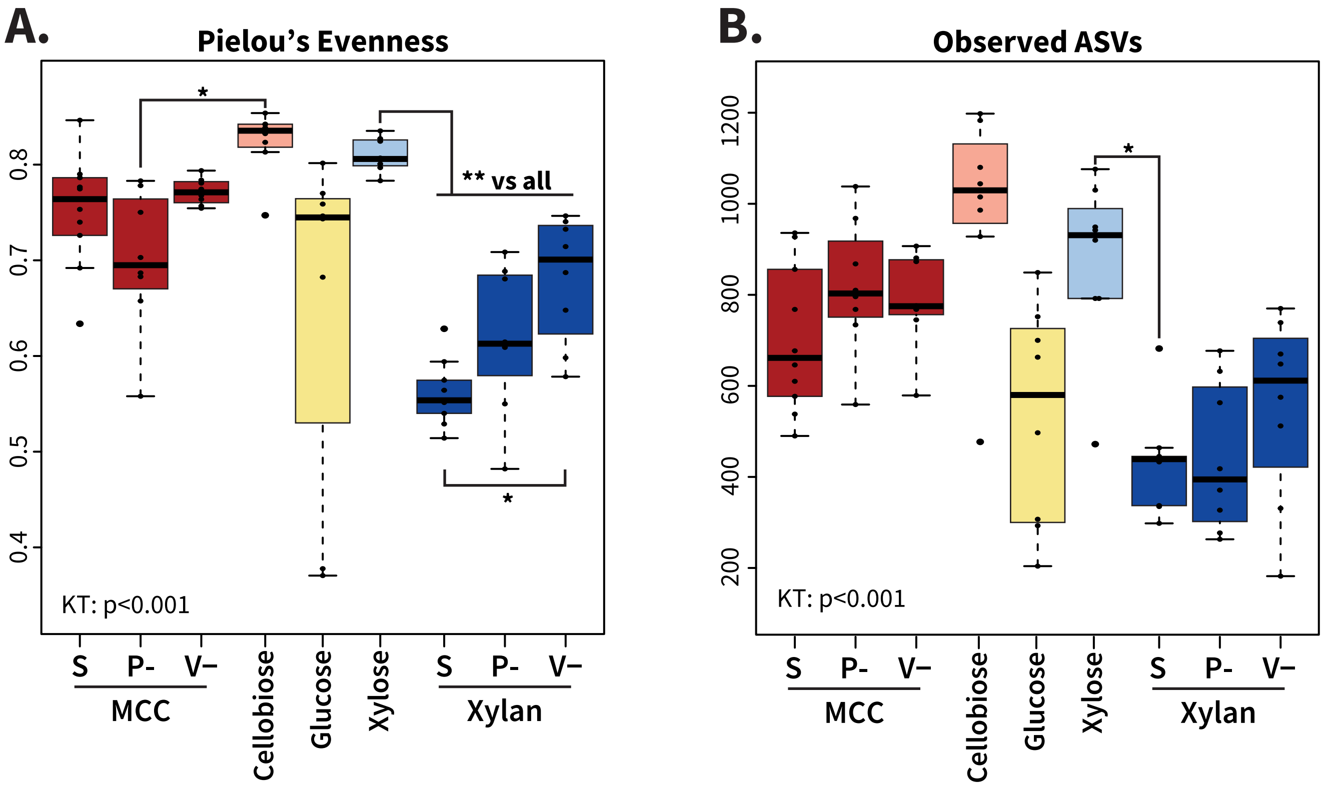


**Supplement 4: Additional alpha diversity measures of standard, deficient, and simple-sugar diets.** Samples were rarefied a constant depth of 12274 sequences for alpha diversity calculations **(A)** Pielou's evenness and **(B)** number of observed ASVs. MCC: microcrystalline cellulose; S: standard diet; P-: protein-deficient; V-: vitamin-deficient. *p < 0.05; **p<0.01

### Supp Fig 5

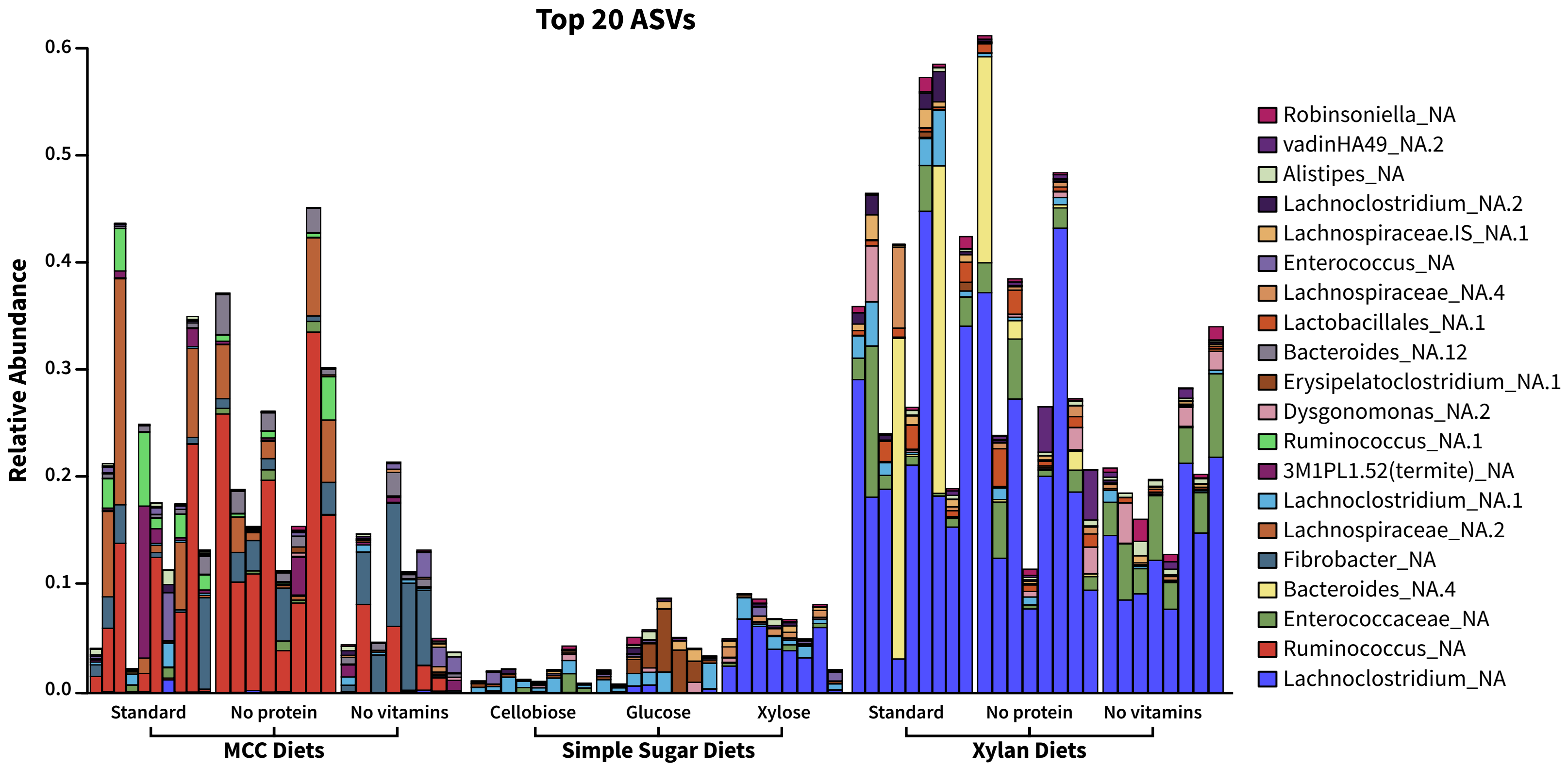


**Supplement 5: Relative abundance of top ASVs in xylan/MCC diet variations.** ASV tables were converted to proportion tables and sorted from most to least abundant ASV across standard, deficient, and simple-sugar variations of the xylan and MCC diets. The top 20 ASVs were included for visualization.

### Supp Fig 6

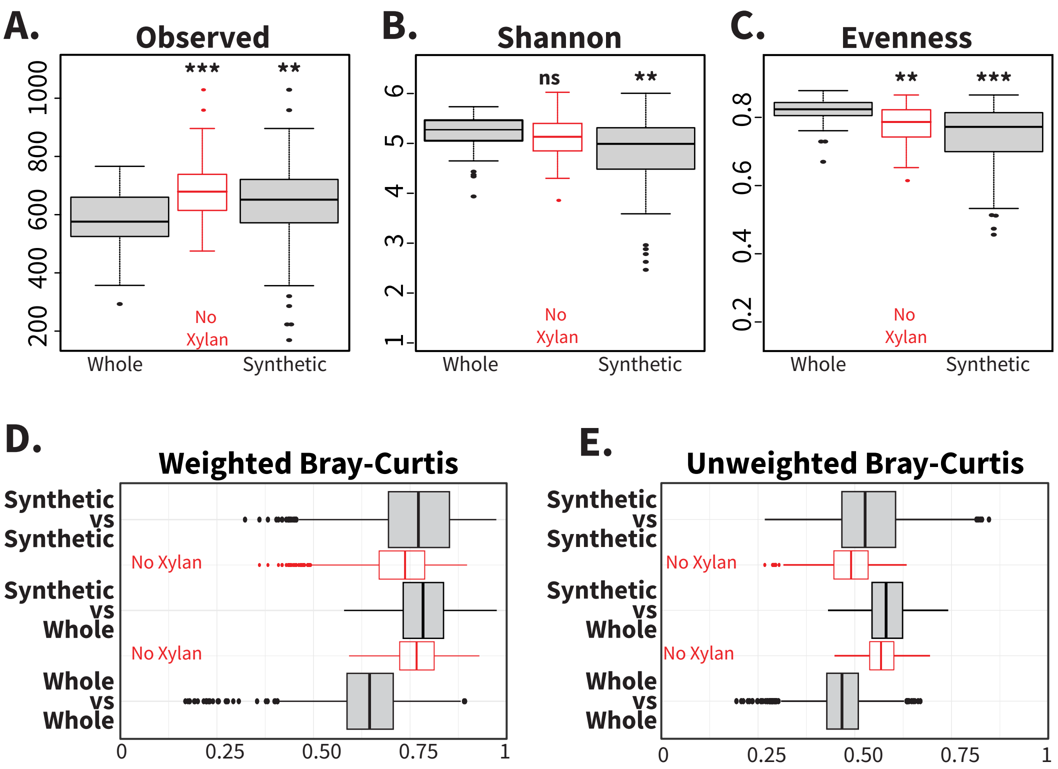


**Supplement 6: Overview of alpha and beta diversity differences between whole food and synthetic diets.** Raw sequence data from Tinker and Ottesen (2016) were reprocessed using the methods in this experiment to generate comparable ASVs. All samples were rarefied to 7924 ASVs for alpha and beta diversity analysis. Boxplots show **(A)** observed ASVs, **(B)** Shannon index, and **(C)** Pielou's evenness for each diet type, with red boxes representing the synthetic diets minus xylan-fed samples. Wilcoxon rank-sum test was used for pairwise statistical analysis between diet types. Beta diversity boxplots of **(D)** weighted and **(E)** unweighted Bray-Curtis dissimilarity, with red boxes representing synthetic diets excluding xylan.  ** =p <0.01; *** =p<0.001, ns =no significance

#### Supp Fig 7

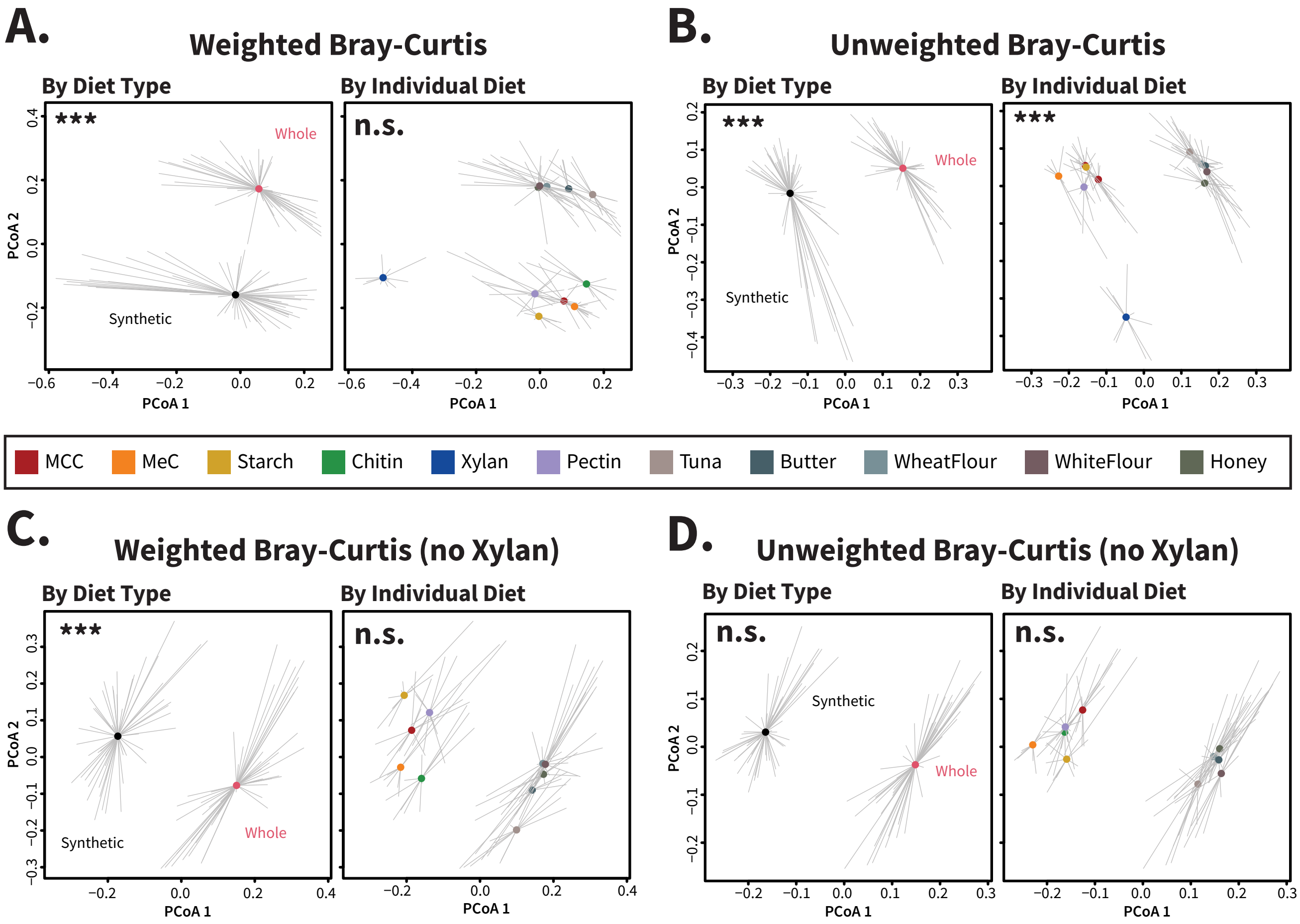


**Supplement 7: Beta dispersion analysis of whole food and synthetic diets, with and without xylan-fed samples.** The dispersion of **(A,C)** weighted and **(B,D)** unweighted Bray-Curtis dissimilarities was assessed for both diet group and individual diets, both **(A,B)** with and **(C,D)** without xylan-fed samples included in analysis. All samples were rarefied to 7924 ASVs for beta dispersion analysis, and ANOVA statistical testing was used to evaluate if dispersion differed between groups/diets. *** =p<0.001, ns =no significance

#### Supp Fig 8

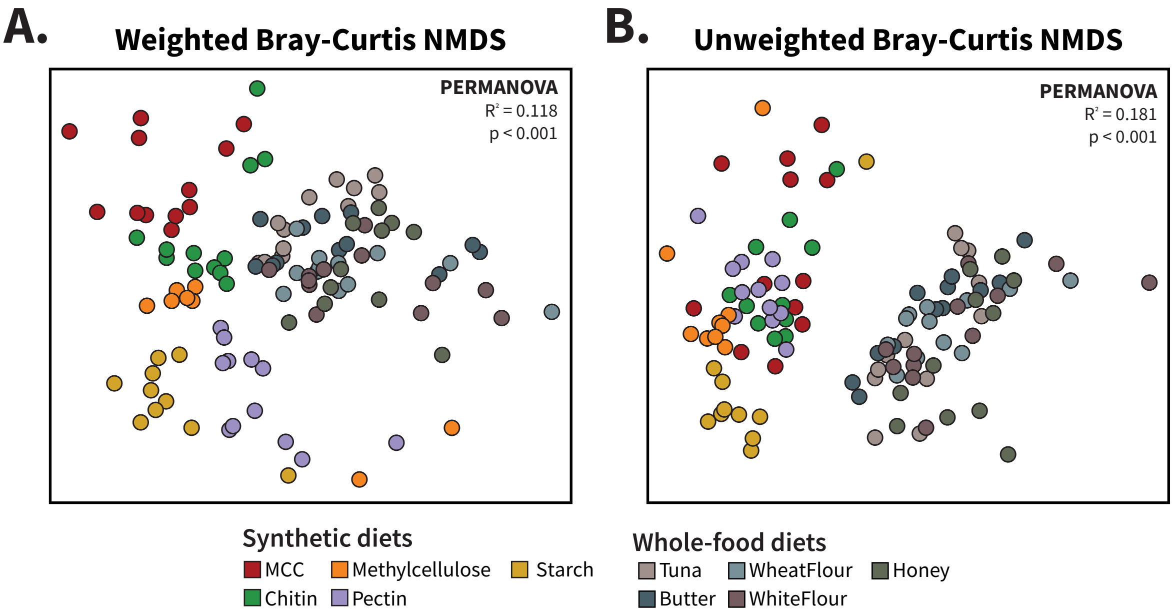


**Supplement 8: Differences in beta diversity between whole food and synthetic diets without xylan-fed samples**. NMDS ordinations of **(A)** weighted and **(B)** unweighted Bray-Curtis distances between cockroach hindgut samples from this study ("Synthetic") and data from Tinker and Ottesen (2016) for cockroaches fed tuna, butter, honey, wheat flour, and white flour ("Whole Food Diets"), excluding xylan-fed samples. Samples were rarefied to 7924 ASVs for beta diversity analysis. $R^2$ and p-values for diet type comparisons were calculated using PERMANOVA.

### Supp Fig 9

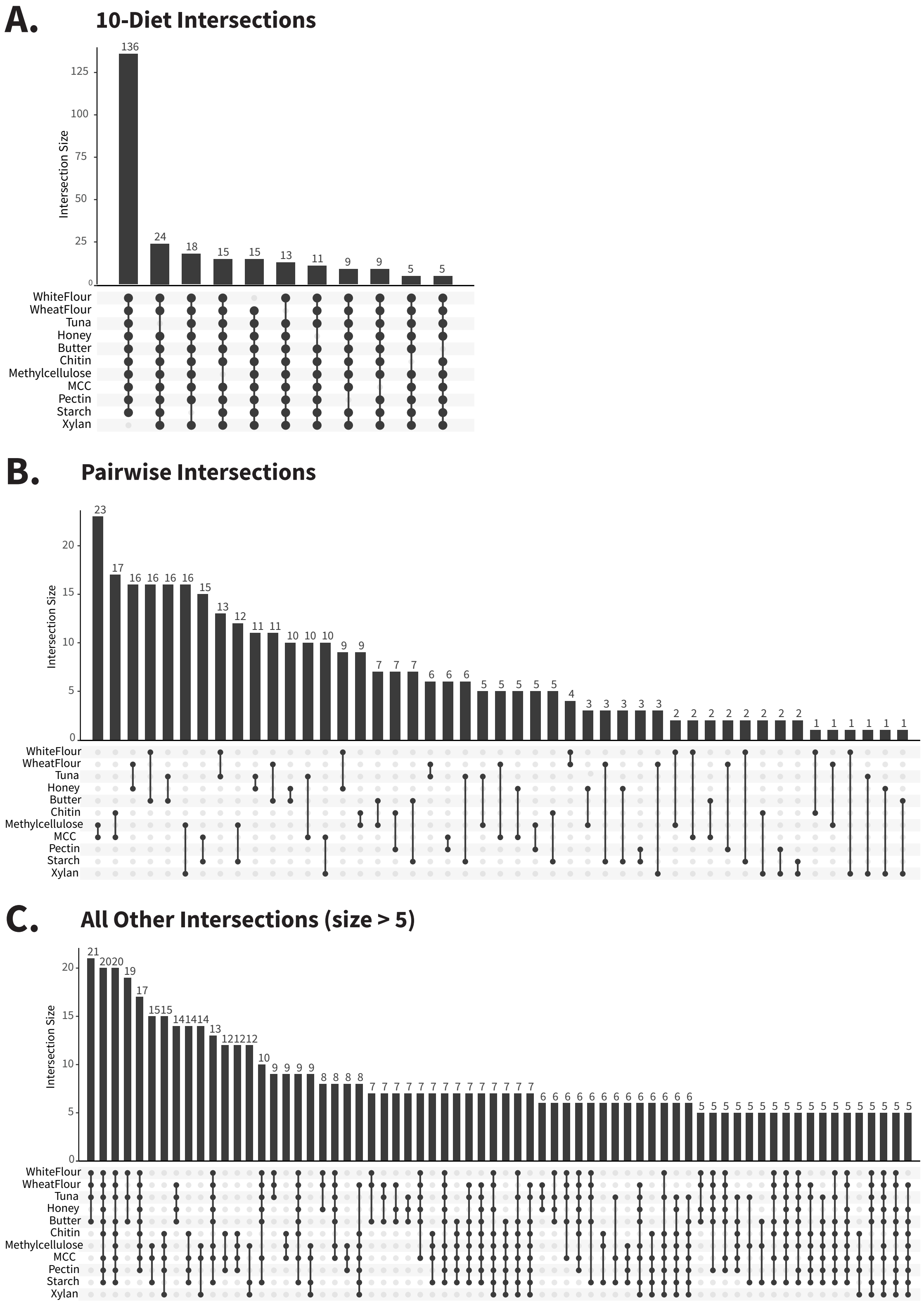


**Supplement 9: Additional UpSet plots.** The rarefied samples (7924 depth) from whole food and synthetic diets were used to generate UpSet plots. **(A)** depicts sets where ASVs are found in all but one of the diets. **(B)** depicts sets found in only two diets. **(C)** contains remaining sets not yet displayed with at least 5 ASVs. Additional sets were not included.

### Supp Fig 10

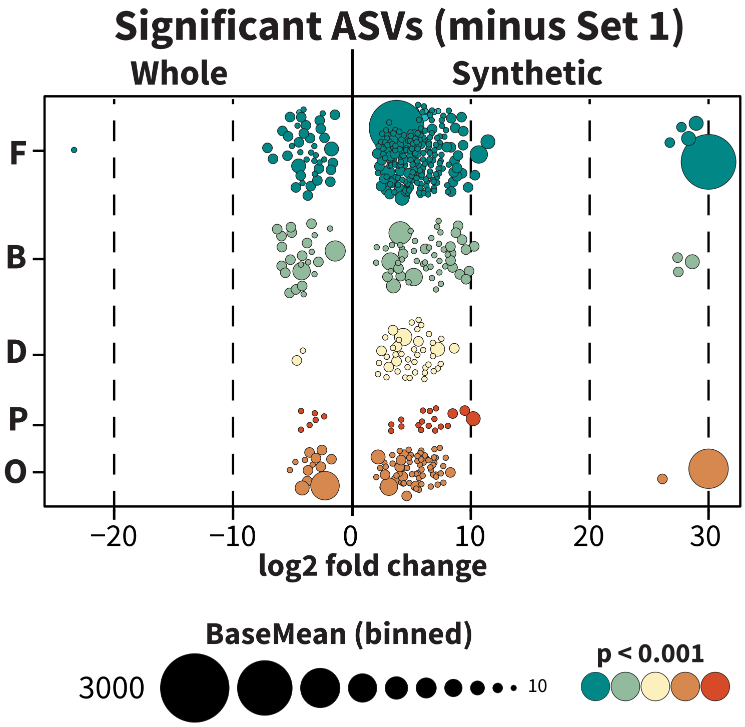


**Supplement 10: Significantly enriched ASVs between whole food and synthetic diets, excluding Set 1 ASVs**. Raw sequence count tables of whole food and synthetic diets were filtered to include ASVs present in at least 5 samples (out of 125 total) and analyzed using DESeq2 with diet type as the design factor. ASVs identified as significant ($p < 0.001$), excluding Set 1, are visualized.

#### Supp Fig 11

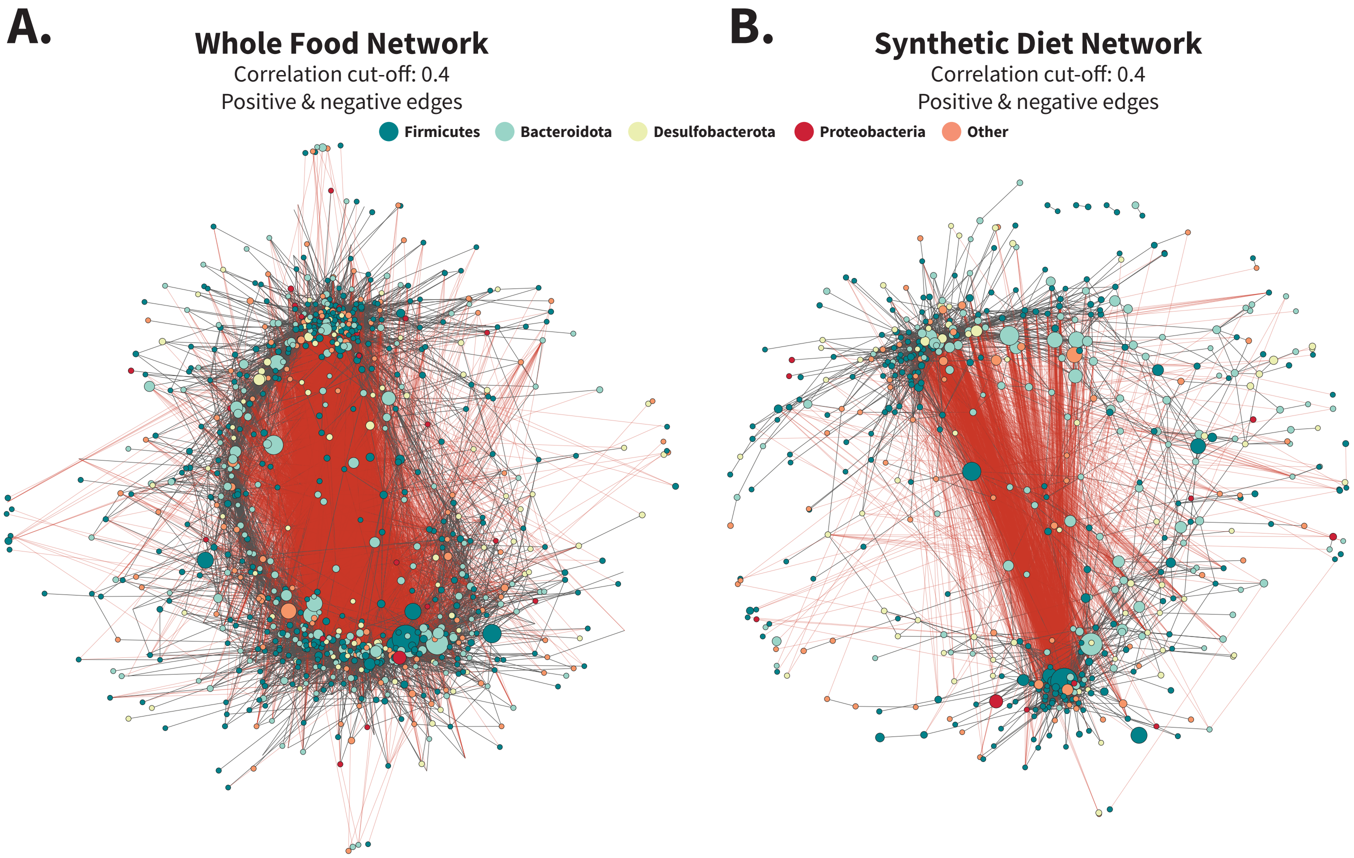


**Supplement 11: Whole food and synthetic networks including negative edges.** Networks were calculated by SparCC from filtered count tables for **(A)** whole food and **(B)** synthetic diets separately to create two distinct networks. Count tables were filtered to include only ASVs present in at least 25% of the samples per diet set, resulting in 976 ASVs for whole food diets and 700 for synthetic. Networks were further pruned to remove edges with absolute values smaller than 0.4 before exporting to Cytoscape. Negative edges were included during initial layout generation with the edge-weighted spring embedded layout method.

### Supp Fig 12

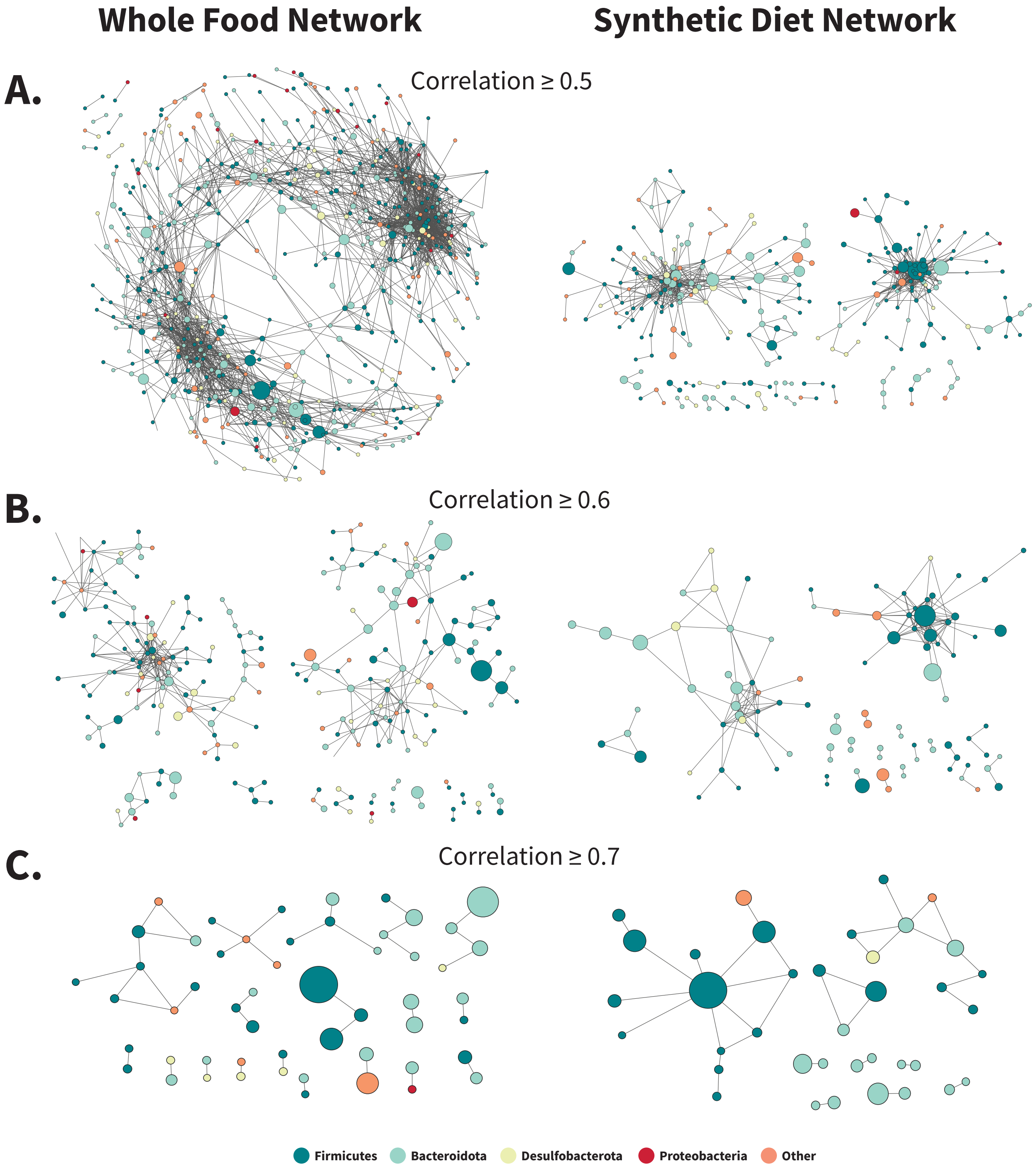


**Supplement 12: Whole food and synthetic networks at higher correlation cut-off levels.** Networks generated for Figures 6 and S11 were filtered in Cytoscape by edge weight to visualize networks at **(A)** 0.5, **(B)** 0.6, and **(C)** 0.7 SparCC correlation values, with isolated nodes removed if they lost all adjacent edges. Data on edges, nodes, and connected components at these cut-off levels are presented in Figure 6C-E.

**Supp Fig 13**

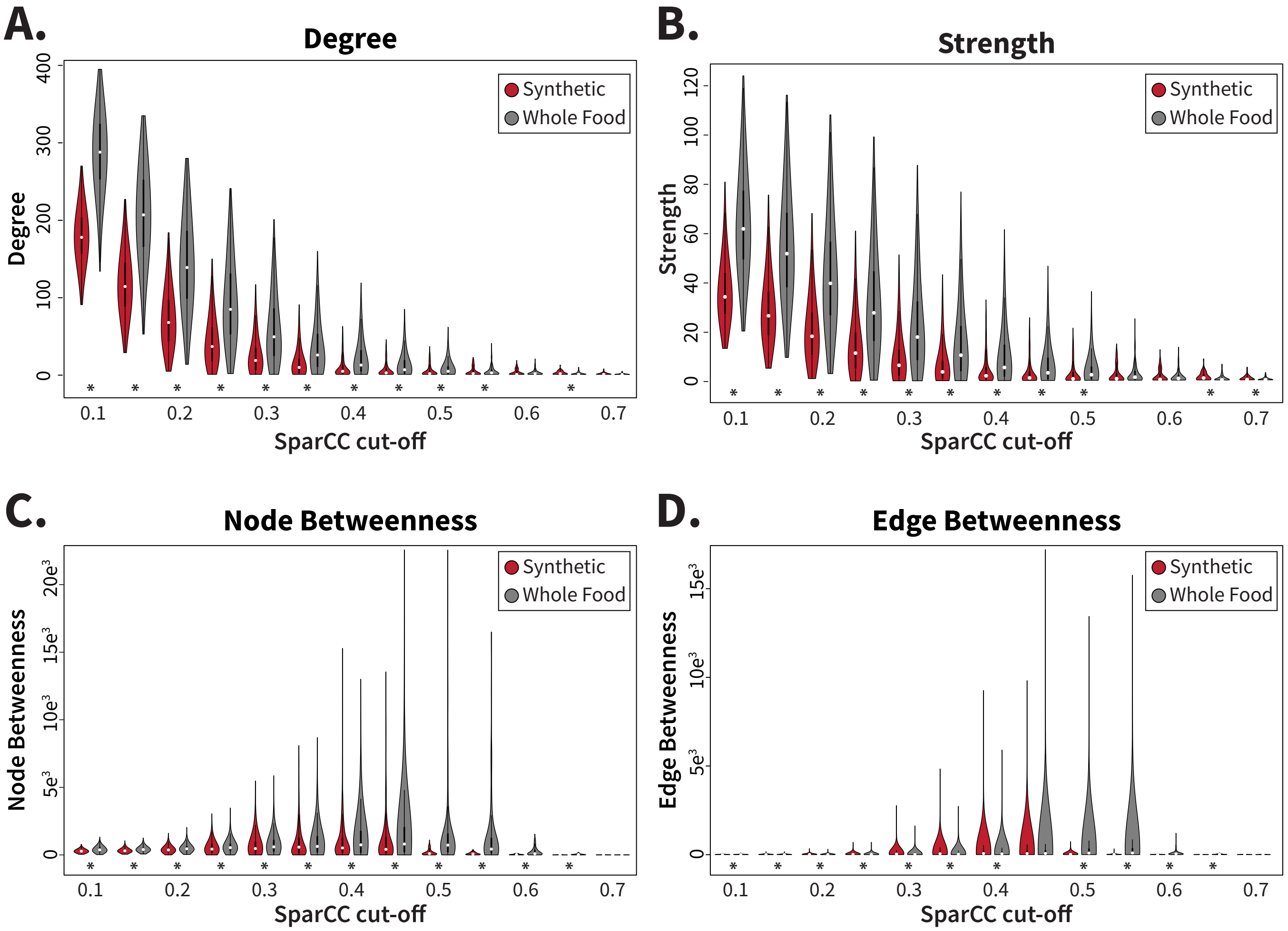


**Supplement 13: Whole food diets encourage more interconnected network formation than synthetic diets.** SparCC networks were generated for whole food and synthetic diet types using raw ASV counts excluding uncommon taxa (< 25% of samples in diet type). The networks were analyzed at increasing levels of positive associations to compare **(A)** degrees per node, **(B)** strength of co-correlations per node, **(C)** node betweenness, and **(D)** edge betweenness. Statistics were calculated using the Wilcoxon rank-sum test. * = $p<0.05$ Red: Synthetic; Grey: Whole Food.
